# Expansion and increase of human pressures on global land ecosystems between 1990 and 2020

**DOI:** 10.64898/2026.04.16.718867

**Authors:** Katharina Ramm, Calum Brown, Almut Arneth, Mark Rounsevell

## Abstract

We present a spatially explicit, global-scale index to assess the effects of the five direct anthropogenic drivers of biodiversity loss identified by the IPBES: land use change, natural resource extraction, climate change, pollution, and invasive alien species. The Biodiversity Pressure Index (BPI) covers 30 years (1990-2020) with an annual time-step and a spatial resolution of 0.1°. We find that the coverage of drivers in available data varies and we highlight the key uncertainties that result from this. Using the best available data, we show that large parts of the terrestrial biosphere (approximately 89%, including Antarctica and Greenland) are under medium or high human pressure and that almost all areas (approximately 96%) have experienced an increase in pressure over the past three decades. The BPI shows varied spatial and temporal patterns across world regions and biomes, but many of these areas are dominated by pressures associated with rising temperatures and trade flows. Tropical and subtropical areas are subject to particularly rapidly-growing pressures, while wetlands consistently show the highest pressure levels across biomes. In revealing these and other patterns, the BPI provides a basis for improved understanding and management of biodiversity impacts in the future.

## 1 Introduction

Anthropogenic pressures have caused a rapid decline in terrestrial biodiversity in recent decades, resulting in extinction rates that are hundreds to a thousand times faster than found in fossil records (De Vos et al., 2015). Although only about 1.4% of species have gone extinct since the industrial revolution (Purvis et al., 2019), an estimated 1 million out of 8 million animal and plant species are now threatened by extinction (Díaz et al., 2019; Purvis et al., 2019), and human pressures continue to increase.

The Intergovernmental science-policy Platform on Biodiversity and Ecosystem Services (IPBES) identifies five main direct drivers of biodiversity loss: land use change, natural resource extraction, climate change, pollution, and invasive alien species (Balvanera et al., 2019). These drivers manifest in diverse threats - habitat degradation, fragmentation, population declines, range shifts, poisoning, and disease. Habitat loss and fragmentation remain the most damaging globally (Davison et al., 2021; Sanderson et al., 2002). Resource extraction *sensu* IPBES includes extraction of living and non-living materials from nature, with localised effects including destruction of habitats and ecosystems (Balvanera et al., 2019; Geist and Lambin, 2002). Natural resource extraction continues to increase, driven by rising populations and per capita consumption, and is distributed more widely through international trade (Schandl et al., 2016). Climate change affects species and ecosystems everywhere, but impacts so far are greatest in higher latitudes, small island nations, low-lying countries, countries with increasingly frequent extreme weather events, and regions prone to wildfires and droughts (Balvanera et al., 2019; IPCC, 2018). Pollution, such as from pesticides and fertilisers, affects groundwater, soil and air quality, with direct and indirect negative impacts on biodiversity (Potter et al., 2010; Sharma et al., 2019; Sud, 2020; Tang et al., 2021). The pollution potential of pesticides is aggravated by their persistence in the environment and consequent long-term impacts (Sharma et al., 2019). Of the other forms of pollution, light pollution has particularly distinct effects, causing irregularities in the natural day-night rhythm and so the metabolism of animals – especially of insects that are attracted to light (Boyes et al., 2021; Hölker et al., 2010). Invasive alien species, mainly introduced through trade and travel, outcompete local fauna and flora, particularly in developed countries with longer histories of trade or larger trade volumes (Roy et al., 2024; Seebens et al., 2017, 2018).

IPBES considers land use change to be the most impactful driver of terrestrial biodiversity loss, followed by resource extraction. Climate change is likely to be the fastest growing threat in the future, with other pressures having more variable negative impacts (Jaureguiberry et al., 2022; Purvis et al., 2019). In any case, all drivers overlap and interact in their biological effects. Addressing biodiversity loss therefore requires integrated strategies that tackle multiple drivers simultaneously and not in isolation (Leclère et al., 2020; Mace et al., 2018).

The Kunming-Montreal Global Biodiversity Framework calls for urgent action to reduce biodiversity pressures and achieve “harmony with nature” by 2050 (Kunming-Montreal Global Biodiversity Framework- 2030 Targets (with Guidance Notes), 2025). This urgency is underscored by the failure to meet most of the preceding Aichi Biodiversity Targets, with some indicators even showing worsening trends that reflect a failure to lessen underpinning pressures on biodiversity (Secretariat of the Convention on Biological Diversity, 2020).

To reduce pressures on biodiversity, we must first understand their extent and magnitude (Bowler et al., 2020). Since these pressures have strong geographic and temporal variation, spatially and temporally explicit datasets are needed to accurately describe them. Increased data availability now makes it possible not only to construct such datasets for each class of pressure in isolation, but also to integrate several pressures into a single index to allow the analysis of combined pressures - and to explore remaining gaps and associated uncertainties in our understanding of the pressures and their effects.

Here we present a Biodiversity Pressure Index (BPI) to quantify the spatial and temporal trends in pressures on biodiversity over 30 years from 1990 to 2020 at high spatial resolution (0.1°). Our aim was to test the feasibility and implications of an index that includes all five of the IPBES direct drivers, despite significant variations in data availability, with resource extraction and invasive species being particularly poorly documented at the global scale. Here, these are represented partially using the most comprehensive data available, along with recommendations for future refinement. The indicators included are land use (types of land use/land cover) and land use transitions (number of transitions between land use/cover types), temperature and precipitation anomalies (i.e. deviations from the long-term mean), pesticide and fertiliser use, light pollution, and trade (imports) as a proxy for potential invasion by non-native species. We also include (limited) data on mining and wildlife trade to represent resource extraction. The land use variable is based on a dataset containing information for both land cover and land management (see methods for more detail on the choice of indicators).

Earlier indices that explored human pressures on biodiversity and ecosystems, such as the Human Footprint (HF) (Venter et al., 2016b), the temporal Human Pressure Index (THPI) (Geldmann et al., 2014) and the Human Modification Index (HMI) (Theobald et al., 2020), have not (with the exception of (Bowler et al., 2020), which includes data on four of the IPBES drivers, but lacks resource extraction) included data on climate change, pollution and/or resource extraction (Geldmann et al., 2014; Venter et al., 2016a). We present our pressure index as an annual timeseries that spans three decades, whereas previous work covered single points in time (Bowler et al., 2020; Sanderson et al., 2002) or shorter or irregular time periods (Geldmann et al., 2014; Mu et al., 2022; Theobald et al., 2020; Venter et al., 2016a) (see Supplementary Table 2 for details). Such a timeseries is necessary to identify how and where pressures change over time, where key data gaps remain, and how policy and conservation efforts might best respond to the changing shape of anthropogenic pressures.

## 2 Results

Globally, human pressures as captured here have increased in extent and magnitude since 1990, with only minor and temporary reductions in the BPI over the period to 2020. The terrestrial area (including Antarctica and Greenland) with a BPI of at least 0.05 (on a scale from 0 to 1; for more information see methods section) increased from 59.3% in 1990 to 88.3% in 2020. Effectively, none (0.46%) of the terrestrial land area was entirely free from human pressure (defined as a BPI of 0) in 2020 (Fig. 1, panels a, b). Overall, between 1990 and 2020, the BPI increased on 90% of terrestrial land, whilst it declined on just 7.7% of land. Around 30% of land showed stable pressure levels over time.

**Figure 1:**
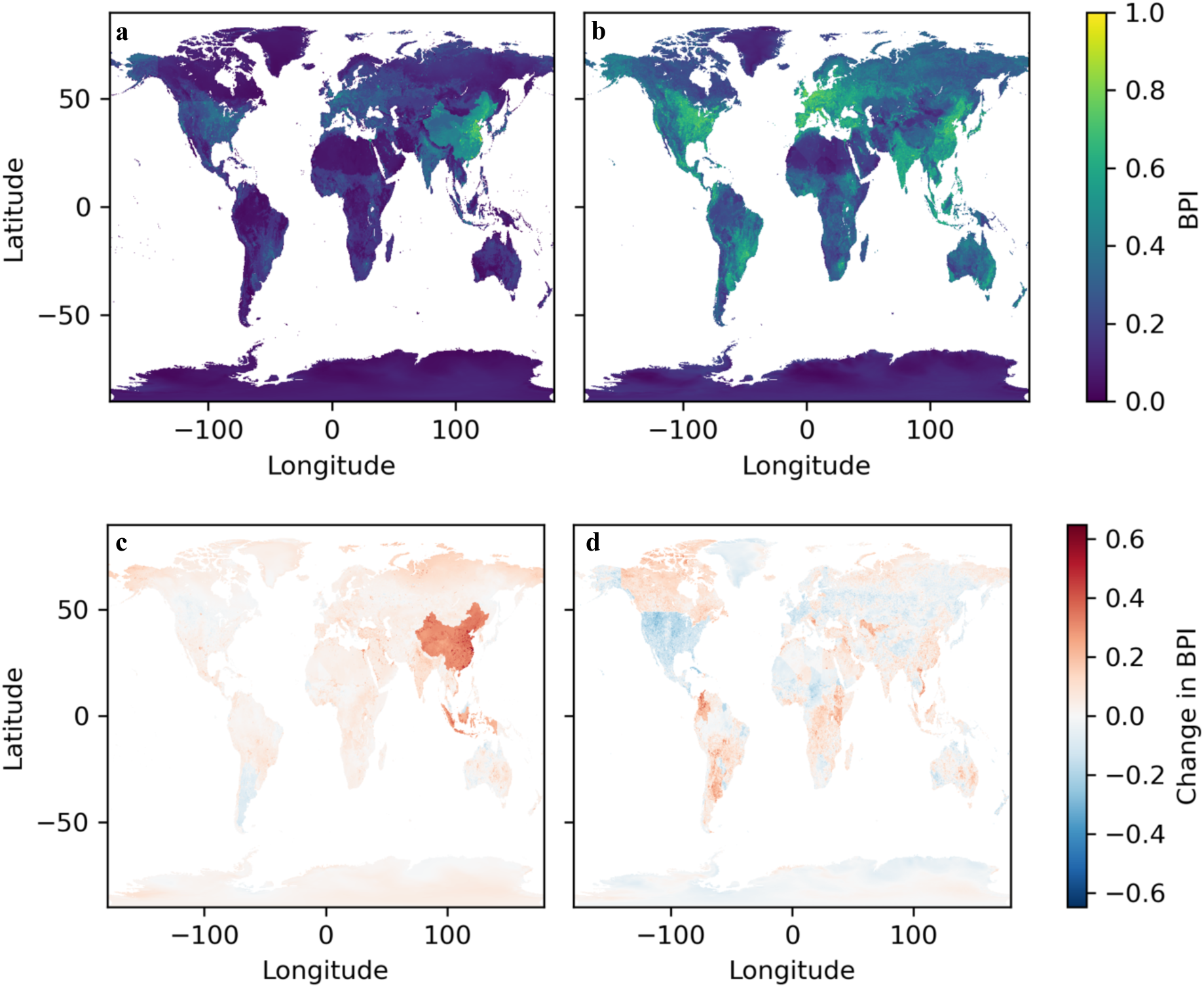
Biodiversity Pressure Index for 2020 (a, b) and change in pressure (c, d). BPI per pixel on a normalised scale from 0 (no pressure) to 1 (very high pressure). Whilst (a, c) show absolute values/changes normalised to [0,1], (b, d) show normalised values/changes of/between percentile ranks, reducing the effect of pressures with large numerical changes that may not relate proportionately to the exerted pressure (see methods for more).

### 2.1 BPI by world regions and biomes

Notable increases have occurred in East Asia & Pacific (which had the highest BPI of 0.24 in 2020, having overtaken South Asia around 2016), particularly in China (driven by trade, temperature, light, and fertiliser use) and Indonesia (wildlife exports, precipitation anomalies - wetter and drier conditions are included in the BPI, land use, fertiliser). Increases also occurred in South Asia (which had the highest BPI of 0.15 in 1990), notably in India (trade, fertiliser, light). Northern latitudes (≥ 55°N) experienced increases due to rising temperatures. Parts of the Middle East & North Africa (which had the lowest BPI in 1990 at 0.05) also showed increases, mainly from light, temperature, and trade. Declines occurred in parts of North America (mainly due to a stop in wildlife exports, regional cooling and reduced land use intensity). Declines also occurred in Latin America & Caribbean, including Argentina (decline in wildlife exports), and parts of Australia (mainly because of land use change) (Fig. 1c, panel c). Europe & Central Asia, together with the Middle East & North Africa, had the lowest BPI in 2020 (∼0.08).

Analysing the BPI in relative terms (using percentile ranking of all BPI values, see methods) minimises the dominance of drivers with large absolute or skewed changes, revealing new hotspots and coldspots of pressure change (Fig. 1, panel d). Aside from influences from climate change as a global phenomenon, regions with more established economies (e.g. North America, Europe & Central Asia, East Asia & Pacific including Japan, Australia) had notably slower increases in pressure than countries with emerging economies (e.g. East Asia & Pacific including China, Latin America & Caribbean including Brazil, Sub-Saharan African states).

Increases in relative pressure (Fig. 1, panel d) in China (mainly from trade, land use transitions and light), while greater in absolute terms (Fig. 1, panel c), were comparable to relative increases in pressure in Brazil (from a mix of pressures, but mainly fertiliser and pesticides), Canada (mainly from trade, land use transitions, temperature), areas in Africa (a mix of pressures, but mainly trade, land use and climate) and the north-western part of South America (from a mix of pressures). Decreases in relative BPI occurred in parts of North America (relative reduction in a mix of pressures, but mainly wildlife trade and temperature), Greenland (mainly because of trade), in boreal areas in Russia (from climate variables, but also from fertiliser), and Sweden (a mix of pressures) as well as Japan (almost all pressures) (Fig. 1, panel d). Pressures in some African regions mainly decreased because of reduced wildlife trade and precipitation anomalies.

All biomes show an increasing trend, with tropical and subtropical dry and humid forests showing the largest increase (0.07) from 0.11 in 1990 to 0.18 in 2020, followed by deserts and xeric shrublands (from 0.06 to 0.11) and wetlands (here limited to a few large tropical and subtropical wetlands, because the data used for biomes from IPBES do not include wetlands from other regions) with an increase from 0.16 in 1990 to 0.22 in 2020. Wetlands were and remain the biome with the highest BPI, above tropical and subtropical dry and humid forests (0.18), temperate grasslands (0.17) and mediterranean forests, woodlands and shrubs (0.17). The lowest values in 2020 were found for the cryosphere (0.01) (Supplementary Fig. 1 and 2 for extents of world regions and biomes).

**Figure 2:**
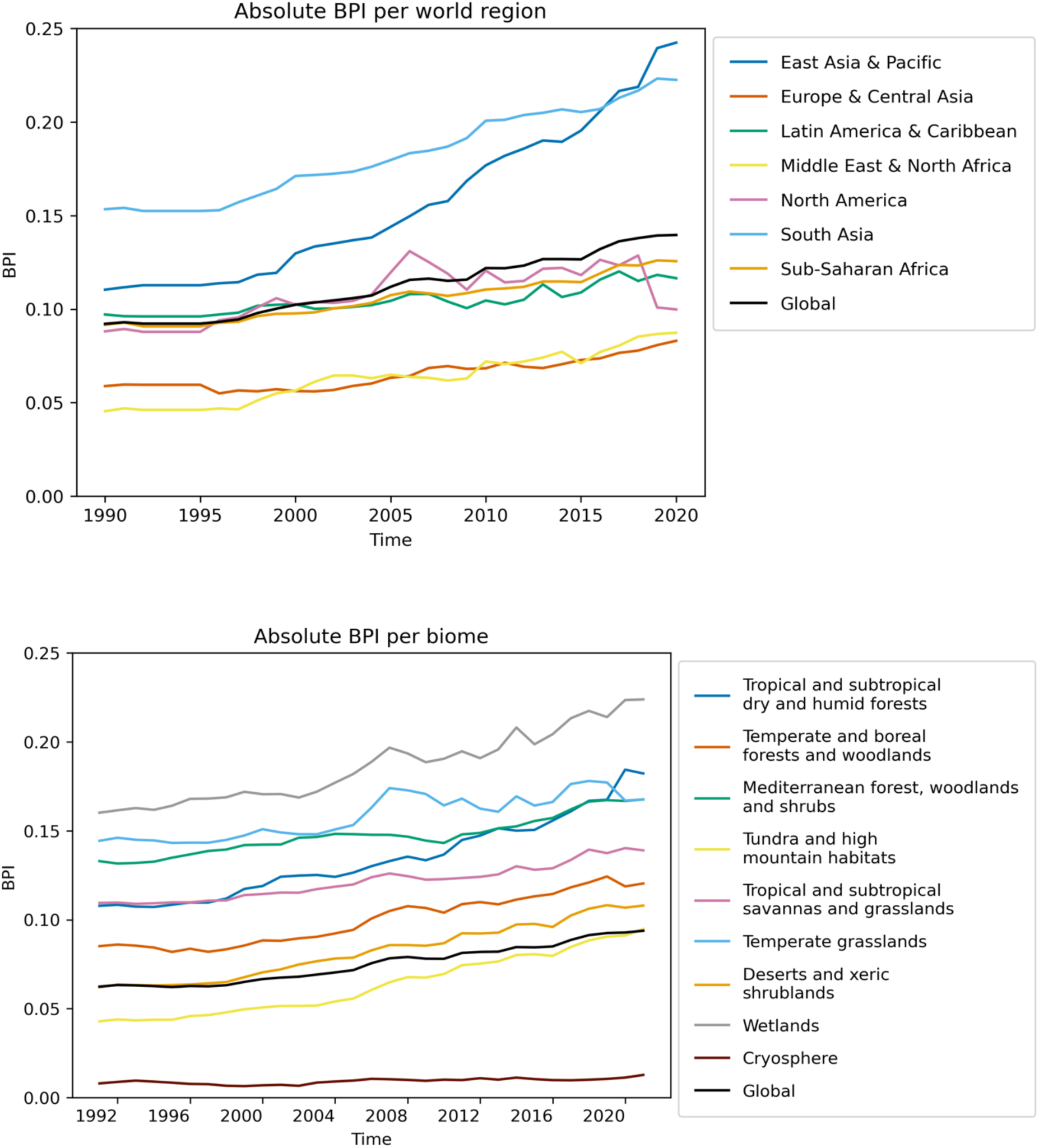
Changes in average BPI from 1990-2020. BPI over time in different world regions (top) and biomes (bottom). Note: wetlands are limited to a few large tropical and subtropical wetlands.

### 2.2 Input variables to the BPI

Overall, absolute pressures have generally increased over the past three decades globally (Fig. 4 and Supplementary Fig. 6), but trends in pressures within world regions and biomes were, as expected, more varied (Fig. 4, Supplementary Fig. 7 and 8). Trade and temperature had the largest increases from 1990-2020, while other pressures increased only moderately or insignificantly or, in the case of wildlife trade, decreased. Land use dominates the global BPI throughout, but warming temperatures have almost reached the same levels in 2020 (Fig. 4). By contrast, when expressing pressures in relative terms, trade emerged more strongly as the pressure with the greatest increase. Wildlife trade also shows high values, as well as the precipitation anomaly and temperature increase.

**Figure 4:**
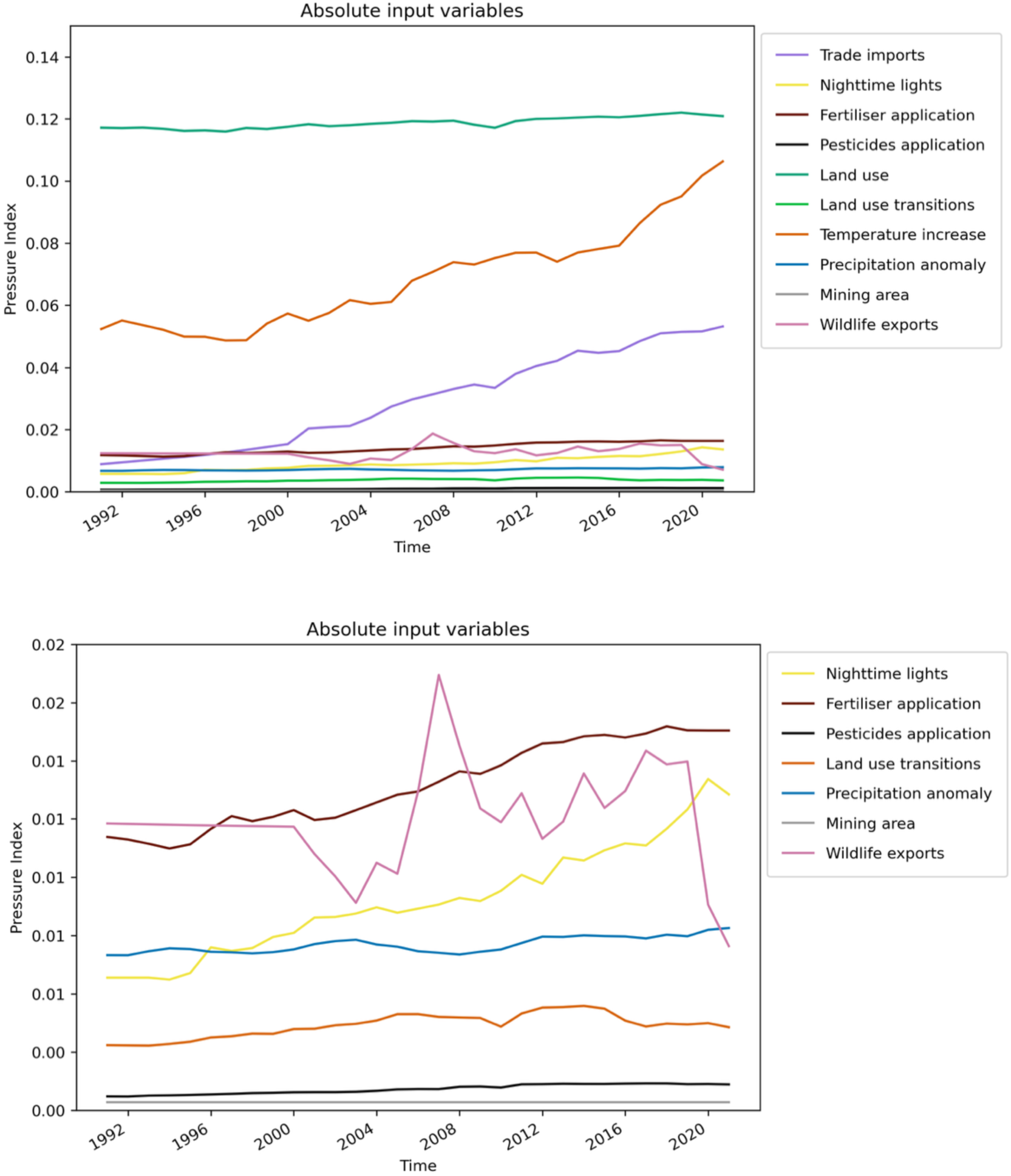
Global mean changes of individual pressures 1990-2020. Each pressure has been normalised to 0-1 and then the yearly means were calculated from the normalised pressures. Top: all input pressures, bottom: without land use, temperature and trade to see more details of the remaining pressures. Further detail on individual inputs can be seen in Supplementary Fig. 5 and 6.

**Figure 5:**
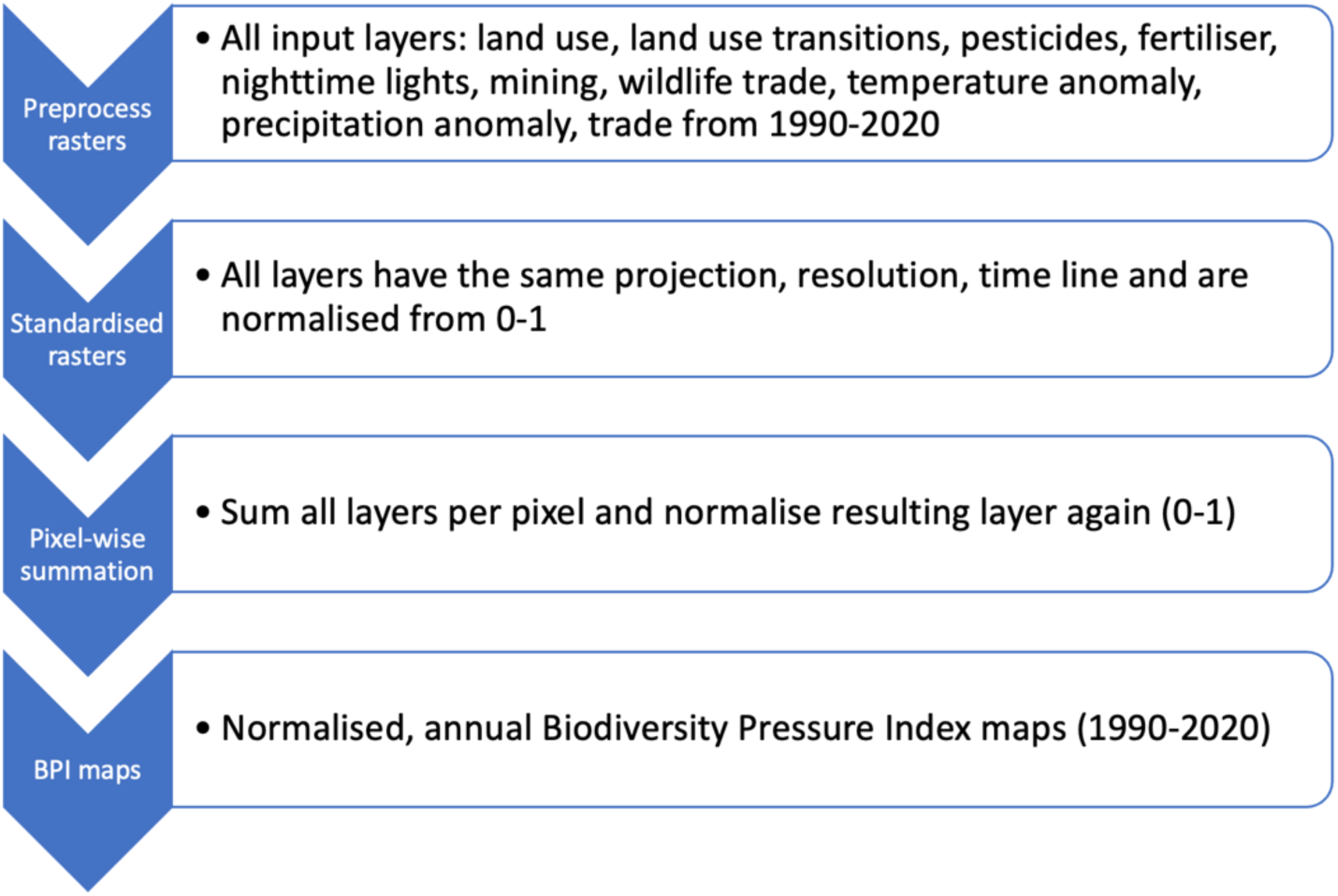
Workflow of the development of the BPI. Showing the step-by-step methodology used to create the BPI.

#### 2.2.1 Dominant pressure

We defined a dominant pressure as one with a normalised value 5% higher than the next-largest pressure (in order to strike a balance between small and uninformative differences and large but rare ones). We find that the climate variables were the most dominant pressures across time and space, as well as the trade variables (Fig. 3). Precipitation anomalies (wetter and drier conditions alike) mainly dominated in tropical and subtropical regions, whilst temperature anomalies (only positive deviations, i.e. increases in temperature) dominated in Antarctica and later also in the Arctic. These were also the pressures with the highest-resolution and best-established datasets, suggesting limited data-related uncertainty at the higher-end of the BPI.

Trade (imports) and wildlife exports also dominated across some extents and periods, but were not strictly comparable because trade values here apply nationally. Thus, values in places such as Alaska and Greenland (being geographically separate from their nation states), as well as in large countries such as Russia, are not always representative of local conditions. Still, the volume of imports made them continually dominant in countries where they were already high in 1990. Land use and land use transitions (with more certain data) dominated parts of Eurasia and in western Africa (e.g., Ghana, Nigeria and neighbouring countries). Regionally, some changes over time were apparent, with warming gradually substituting trade as the dominant pressure in northern latitudes (especially in Greenland and Russia), and land use transitions increasingly dominant in Europe and Australia, mainly in the last decade (2010-2020) (Supplementary Fig. 4).

#### 2.2.2 Land use and land use change

Land use pressure was stable in all world regions and plays a major role in South Asia and in East Asia & Pacific, Sub-Saharan Africa and Latin America & Caribbean; also affecting wetlands and the Mediterranean forest, woodlands and shrubs and other biomes. Land use transitions (number of changes between land uses) were most prominent in South Asia and East Asia & Pacific and in the Mediterranean forest, woodlands and shrubs and least prominent in the Middle East & North Africa and the Cryosphere. Overall, land use transitions were on average relatively stable, but frequency of transitions increased in some and decreased in other regions with some increases and some decreases in different regions.

#### 2.2.3 Resource extraction

Wildlife exports were not so important and less varied in South Asia, Europe & Central Asia and Middle East & North Africa and most prominent in North America, with a steep drop in 2019 (potentially an artefact of the data). Mining is a stable variable, so no changes between the years could be derived. Nevertheless, mining plays a large role in Latin America & Caribbean and East Asia & Pacific as well as in the tropical and subtropical dry and humid forest biomes and in Mediterranean forests, woodlands and shrubs.

#### 2.2.4 Environmental pollution

Fertiliser played a large role in South Asia and to a lesser extent in East Asia & Pacific, increasing most in wetland biomes as well as in the tropical and subtropical dry and humid forest biome. Pesticide use increased most strongly in Latin America & the Caribbean, in East Asia & Pacific, and in tropical and subtropical dry and humid forest and savannas and in the temperate grassland biomes. Night time lights increased substantially in the Middle East & North Africa and a little less in South Asia, while increasing most in the Mediterranean forest, woodlands and shrubs as well as in the wetland biomes.

#### 2.2.5 Climate change

Temperatures warmed in all world regions, with the largest increases in Middle East & North Africa and, in terms of biomes, in deserts and xeric shrublands. Precipitation both increased and declined, and anomalies were thus positive and negative with overall no clear trend. Precipitation nevertheless plays a substantial role in tropical and subtropical grasslands and forests and in wetlands, as well as in Latin America & Caribbean and other regions.

#### 2.2.6 Alien invasive species

Trade played and plays a relatively large role in North America but did not show a large increase in the last three decades. Trade on the other hand increased sharply from 2000 onwards in East Asia & Pacific and South Asia, while also increasing substantially across most biomes.

## 3 Discussion

### 3.1 Global findings and differences to other indices on a global scale

Using available global data to describe the five main anthropogenic drivers of biodiversity loss over the past 30 years, we find that human pressures on biodiversity increased across all world regions and biomes, with 90% of the terrestrial land area experiencing an increase in pressure (mainly due to trade and temperature). The magnitudes of these changes are well-represented by available data, but their spatial occurrence remains uncertain, largely due to the cryptic nature of trade impacts. Only relatively small areas experienced few, small, or even no pressures in 2020. The areas less-affected by anthropogenic pressures tended to be in the Arctic and other environments unsuitable for agriculture and other impactful human activities. The general global increase and broad patterns of pressures in BPI agree with those described in the regional assessment reports by (IPBES, 2018c, b, d, a) and the studies by (Geldmann et al., 2014; Theobald et al., 2020; Venter et al., 2016b) and thus highlight the need for urgent protection of the environment in light of rising pressures.

We compared some of our results to findings from other studies with similar approaches (Table 4). We find 1.2 times more land area affected in the BPI compared to a widely used earlier approach, the Human Footprint (HF) maps (Venter et al., 2016b). However, the BPI includes Greenland and Antarctica, whereas the HF does not. In terms of global mean pressures (relative to their potential maximum pressure), the HF maps and the BPI show similar values in 2009 (∼12% and 8% respectively), while the mean value of the Human Modification Index (HMI) (Theobald et al., 2020), another similar index that excludes Antarctica, is the same as the BPI (9%) in 2015. These mean values are low due to the large skew in the data, with exceptionally high values recorded in small areas and large parts of the globe with low to medium pressure.

Global rates of change also differ. The HMI increased by about 15% from 1990 to 2015, whilst the BPI increased by 36% over the same period. The Temporal Human Pressure Index (THPI) (Geldmann et al., 2014) showed that 64% of all terrestrial areas (without Antarctica) experienced an increase in anthropogenic pressure from the early 1990s to the late 2000s (about 15 years). Over the same time period, we found an increase in pressure across 76% of the land areas in the BPI. The higher values for extent and rate of increase in the BPI are due to the inclusion of additional pressures, with geographically dispersed effects.

Relatively intact habitats in the HF maps (i.e., areas with initial pressure less than 4 out of 50), (Venter et al., 2016b) declined in area by 9.3% from 1993 to 2009 (Venter et al., 2016b). By comparison, if we assume that areas with an initial BPI of less than 0.05 out of 1 to be mostly intact, we calculate a decrease in intact area of 15% over the same time period. Another study derived from the HF maps estimated that in 2009, ca. 23% of terrestrial land (excluding Antarctica and Greenland) was still ‘wild’, based on HF values of less than 2% of the maximum (Watson et al., 2016). According to our BPI, only a negligible 0.5% of terrestrial land was wild in 2009, using the same threshold of 2%. Thus, in our study, wild areas were about 46 times smaller, but both the HP and the BPI locate these areas in similar parts of the world such as the Canadian and Siberian boreal/tundra regions, the Sahara, parts of the Amazon Forest, Indonesia and the Australian outback and parts of western China (Watson et al., 2016). These differences again underline the pervasive nature of climate change in extending human impacts far into otherwise unaffected areas. The HP approach did not include climate variables, whereas in the BPI, climate variables (especially temperature) were shown to be an important driver of increasing pressure.

**Table. 1:** Comparison of BPI with other pressure indices. Values are based on publications of the respective indices. More detail on the data included, temporal coverage and spatial resolution can be found in the Supplement.

| Index | Mean pressure (proportion of maximum) | Increase in global mean pressure | Increase in area affected by pressure | Land area affected | Decrease in intact land | Wild area left | IPBES drivers included |
| --- | --- | --- | --- | --- | --- | --- | --- |
| Human Footprint (Venter et al., 2016a) | 12.32% in 2009 (max. pressure value = 50) | x | x | 75% in 2009 | 9.3% from 1990-2009 | 23% in 2009 | Land use change, alien invasive species |
| Human Modification Index (Theobald et al., 2020) | 9% in 2015 (max. pressure value = 1) | Percent change = 15% from 1990-2015 | x | x | x | x | Land use change, resource extraction, pollution (not from agriculture), alien invasive species |
| Temporal Human Pressure Index (Geldmann et al., 2014) | x | x | 64% ~1990-2009 | x | x | x | Land use change, alien invasive species |
| BPI | 8% in 2009<br>9% in 2015 (max. pressure value = 1) | Percent change = 36% from 1990-2015 | 76% from 1990-2009 | 90% in 2009 | 15% from 1990-2009 | 0.5% in 2009 | Land use change, resource extraction, pollution, climate change, alien invasive species |

The BPI generally shows higher pressure levels than previous indices, likely because it includes all five IPBES direct drivers, which earlier studies did not. Climate change, especially rising temperatures, plays a major role in shaping BPI trends and this affects even remote areas. Our analysis also covers Antarctica and Greenland, which were excluded or only partially covered in past work (Bowler et al., 2020; Geldmann et al., 2014; Theobald et al., 2020; Venter et al., 2016a). Furthermore, trade (import) values (as a surrogate for the potential invasion of alien species) and wildlife exports in the BPI apply at the national level, because it is not yet possible to trace or anticipate the spread of the many pests, pathogens and other potentially invasive species that trade might facilitate within countries. While we feel it is important to include these data we acknowledge that without geospatially explicit information the BPI is likely to overestimate the rate at which invasive species could affect more remote areas, or countries whose imports are less likely to host invasive species (Chapman et al., 2017; Seebens et al., 2018). It may also obscure potentially intense localised pressures from trade where traded materials or invasive species are concentrated. As such, the pressures of trade may be over- or under-estimated, depending on the nature of the processes involved.

### 3.2 Regional findings

We found that East Asia & Pacific had the highest absolute regional value of BPI in 2020, mainly caused by land use, fertiliser, trade, wildlife exports and temperature. The temperate and tropical forests of China and Southeast Asia were also highlighted previously as having high pressures by (Venter et al., 2016b). Asia (together with Oceania and Europe), saw the greatest human modification between 1990 and 2015 according to the study of (Theobald et al., 2020). This aligns with the observed rapid loss of biodiversity in tropical Asian regions (Ceballos et al., 2020), often attributed to replacing natural forest with tree crops, along with hunting and wildlife trade, mining (of e.g., limestone for cement), wetland drainage and associated peat fires (Hughes, 2017). Rapid population and economic growth have also been linked to recent declines in biodiversity in China (Ma et al., 2022).

Europe & Central Asia and the Middle East & North Africa showed the lowest absolute regional value of BPI in 2020. Other studies also found that the desert regions of Africa, for instance, had minimal pressures (Geldmann et al., 2014; Sanderson et al., 2002; Venter et al., 2016b). We observed notable BPI increases in areas such as the Mediterranean coast of Morocco and Algeria, the Nile and its delta, and parts of Saudi Arabia. However, these smaller high BPI zones were outweighed by extensive low BPI arid regions. Previous studies suggest that rapid population growth, urbanization, infrastructure development, desertification, overexploitation, and invasive species may pose growing threats to biodiversity in these regions, which the BPI – despite its data limitations – seems to be picking up (Ayyad, 2003; Darkoh, 2003; Jowkar et al., 2016).

Among biomes, tropical and subtropical wetlands, dry and humid forests, Mediterranean forests, woodlands and shrubs, and temperate grasslands had the highest BPI in 2020 - echoing findings of heavy human alteration in these biomes (Williams et al., 2020). Wetlands, often drained for agriculture and impacted by pollution, declined globally by 31% between 1970–2008 (Dixon et al., 2016), with most losses in tropical and subtropical areas (Balvanera et al., 2019; UNEP, 2016), which are the regions covered by the BPI analysis. With a pixel size of around 10km at the equator, the BPI probably underestimated pressure on smaller wetlands that were not considered in the analysis. Only an estimated 0.81% of all temperate grasslands remain wild (Williams et al., 2020), and regions that would naturally be grassland have undergone the highest modification by humans outside the tropical dry forests biome (Jacobson et al., 2019). Temperate grasslands are often converted to agricultural land where soil is productive, or used as pasture where it is not. Consequently almost no grasslands in Europe or North America remain untouched (Baumann et al., 2020; Ceballos et al., 2010; Gibson, 2009; WWF, 2023). Mediterranean areas are very negatively influenced by increasing temperatures (Miranda et al., 2023) and historic land use (Gauquelin et al., 2018). The BPI results resonate with other findings that show biodiversity being especially vulnerable to these changes in that area (Newbold et al., 2020).

### 3.3 Findings from individual pressures

Whilst the total BPI has increased, trends in the individual pressures have been variable over the last three decades and in different regions. Still, there are some common patterns: (1) Both trade and temperature have increased strongly and now, together with land use, dominate the BPI (see Fig. 4) in absolute terms. Trade grew steadily, with a small drop during the global financial crisis. Global trade has long been seen as the main pathway for invasive alien species (Hulme, 2009, 2021) that have direct and indirect adverse effects on landscapes, ecosystem services and native species (Epanchin-Niell et al., 2021). (2) Temperature and trade are the fastest growing pressures and climate warming is expected to reach or even supersede land use in the future as the pressure with the highest contributions to the BPI globally. A warming climate and a higher frequency of extreme weather events (IPCC, 2023) is known to contribute to change and potential loss of habitats as well as to changes in species traits, behaviour, and extinction risks (Habibullah et al., 2022; Jetz et al., 2007; Nunez et al., 2019; Weiskopf et al., 2020). The increasing importance of temperature, captured by the index, implies therefore substantial impacts on biodiversity. For example, Nunez et al, (2019) found that if temperatures increase up to 2°C above pre-industrial levels, species (plants and animals together) could be reduced by about 14% in terrestrial ecosystems and species lose 35% of their climatically suitable habitat. (3) Even though land use (in terms of global pressure levels) did not change markedly over the study period, it remained the driver with the highest normalised values globally until 2013 and the second highest thereafter. Urbanisation and agricultural expansion affect local biodiversity and lead to an increase in BPI. Urban areas increased particularly in Asia, Europe and North America, where for example especially small urban areas have expanded markedly (Zhao et al., 2022). Cropland expansion occurred widely, and offset any cropland losses (Winkler et al., 2021). Net levels of land use pressures are thus relatively stable.

### 3.4 Methodological findings

The BPI is a new spatial index to explore annual human pressure related to biodiversity loss, here calculated for a 30-year period at a spatial resolution of 0.1° and annual time step. The content and construction of the index is flexible, with potentially substitutable datasets to describe individual pressures, and no fixed assumptions about their numerical combination or weighting. This means that the index can be tailored to particular contexts and objectives Importantly, it can be easily updated to accommodate improved data sources.

The input variables to the BPI (land use, land use transitions, mining, wildlife trade, pesticides and fertiliser pollution, night time lights, temperature and precipitation anomalies, trade) were chosen to represent all IPBES-defined direct drivers (Balvanera et al., 2019) on a global scale using open data. It is only now that it is possible to represent all IPBES drivers with the available data and the BPI is partly an experimental metric to assess the utility of this representation. As such, the data used in this study have limitations in terms of spatial and temporal resolution, uncertainties and potential inconsistencies, and are used here in lieu of more detailed data on pressures within each of the classes (see the Supplementary, table 1 and 3 for more information on the input data of this study). Data resolution is an important limitation when calculating the BPI, and better resolved data would markedly improve the capacity to identify regional differences and specificities. Comparing pressures measured in the current way with those measured using detailed data, where and when available, would provide valuable information for interpretation of pressure indices.

Data availability also limits appropriate representation of the IPBES drivers. Even though this study includes all of the main classes of direct drivers, it does not account for all potential sub-classes and might therefore under- or mis-estimate the overall pressure (Williams et al., 2020). Additionally, some variables, such as fertiliser or lights, give more direct representations of pressures (land use intensity, pollution), compared to e.g. (wildlife) trade, which only acts as a proxy (for resource extraction and alien invasive species). Moreover, pressures usually do not act alone (Bowler et al., 2020), and it is not clear whether or when their impact is additive (as currently represented in the BPI) or interactive (Côté et al., 2016). These forms of interactions also vary among environments and species. Furthermore, the nature of the datasets did not allow us to look at freshwater systems and hydrological basins, which have the capacity to increase the area of impact from e.g. dissolved pollutants beyond the area of their application, as well as being directly impacted themselves by pollution (Cable et al., 2017; Maggi et al., 2023).

Importantly, the BPI is built to show human pressures, and not their realised impacts on biodiversity. The translation from such an index into realised impacts is a distinct and substantial task, requiring contextual information and interpretation (Geldmann et al., 2014; Halpern and Fujita, 2013; Williams et al., 2020). This makes the BPI more suited to analyses at large scales and in the context of science-policy processes than for local-scale management. Pressures - individually and combined - have highly variable effects on different species, also depending on ecological and societal circumstances (Foden et al., 2013; Newbold, 2018; Pereira et al., 2012). The BPI cannot therefore be interpreted as a direct measure of realised human impact on biodiversity, but can be used to identify places at high risk of negative impacts, and how those risks have changed over time.

It is clear from these (and other) findings that the *potential* for substantial negative impacts on natural systems has increased rapidly and expanded over recent years. We find no signal of a concerted slowing or reversal of these trends, even following global agreements on reducing pressures and reversing the loss of biodiversity. The BPI can help to track changes in pressure over time and thus help to formulate better tailored nature conservation measures. For example, the BPI would be a suitable approach to explore human pressures across protected areas, and how pressures vary between different protection levels, size, age and other attributes of the protected areas themselves. Whilst the historic BPI can help to reveal success and failure of nature conservation measures in the past, a BPI based on future projections would also show where pressures could change under different scenarios. However, further work would be required to develop future scenarios of the BPI, including projecting forward the individual variables that make up the index. Understanding why and where human pressure is growing (not only within, but also outside of protected areas) in the past and the future is essential in reducing human pressures or their impacts on biodiversity. This also requires an exploration of how pressures act in combination. National biodiversity strategies are directly linked to the CBD (Convention of Biological Diversity) and to findings from IPBES. A BPI based on IPBES drivers could therefore directly support research and global conservation efforts.

## 4 Data and methods

In the development of the BPI we built on previously proposed methods (Bowler et al., 2020; Geldmann et al., 2014; Sanderson et al., 2002; Venter et al., 2016a). The BPI was calculated as an annual index for the years 1990-2020 with a spatial resolution of 0.1° from various global-scale indicators. These indicators were selected to represent land use, natural resource extraction, climate change, pollution, and invasive alien species, reflecting the main direct drivers of global biodiversity change defined by IPBES (for more detail see table S1). An overview of the global indicator datasets is provided in table 2 in the Supplementary. The input variables were pre-processed (see sections below) and summed per pixel with equal weighting for all variables resulting in pressure maps with values ranging from 0 (= no or low pressure) to 1 (= high pressure).

The data preprocessing and the calculation of the BPI was conducted with R 1.4.1103 and Python 3.10. All data were processed in a geographical lat/lon projection with WGS84.

### 4.1 Approach to data selection

Pressures, also called direct drivers by IPBES, refer to direct, human-induced (anthropogenic) pressures on biodiversity. We did not consider purely natural or indirect drivers. In order to be considered for the analysis, indicator datasets had to fulfill certain requirements in being: a) global terrestrial data (freshwater was not considered individually and exceptions were made for datasets not including polar regions) , b) spatially-explicit at high resolution, c) time-series including (if possible all) the years from 1990 and 2020, d) open access and of high quality (e.g. peer-reviewed), e) methods that were consistent and comparable for datasets with several measurements. The aim was to represent the direct drivers of biodiversity loss identified by IPBES as well as possible. This was challenging due to the scarcity of appropriate data, with most being not spatially explicit, representing only a small region, or only describing one point in time. Where timeseries existed, it was necessary in some cases to deal with gaps in their temporal coverage. In total, nine datasets (see also SI, table 2) were retained for the analysis from the list of possibly useful datasets, after checking against the selection criteria. All but one (land use) datasets were metric and all but one (mining) were available as a time-series.

The selected variables represent various potential impacts on biodiversity directly or serve as surrogates for pressures identified by IPBES. Correlation between variables exists because of real-world correlation (and compounding) of pressures, but can also occur where datasets share input data. A study by Bowler et al. (2020), for instance, explored common combinations of pressures and found that some pressures are more likely to appear together, i.e. are more correlated (termed anthropogenic threat complexes by the authors). In our case, nighttime lights are used to indicate light pollution, but also indicate the presence of infrastructure. Similarly, agricultural land conversion and application of chemicals can affect biodiversity in multiple ways, while generally occurring in the same place. However, to minimise the extent of artificial correlation, we have only used distinct datasets (if available) across pressures so that land use data, for example, are not repeated in the calculation of fertiliser application. Hence, we have deliberately selected alternate datasets, where possible, to represent the direct drivers. While this is likely to introduce some inconsistencies between pressures, it avoids excessive replication of single datasets.

### 4.2 Land use and land use transitions

The HILDA+ land use reconstruction is a global product covering the years 1960-2020 showing major land use classes on a 1 km grid. It is based on a combination of satellite and statistical data (Winkler et al., 2021). Three different datasets from HILDA+ were used in the BPI:

The land use classes showing a land use for each pixel. We assigned each land use class a pressure value (from 0 to 1) depending on its potential impact on biodiversity following the approach of Sanderson et al. (Sanderson et al., 2002; Venter et al., 2016a). Urban environment received a score of 1.0, cropland 0.7, pasture 0.4, managed forest from 0.1 to 0.5, unmanaged forest, scrubland, other land and water 0.0. This data set was used to represent existence and expansion (or decrease) of certain land use types e.g. agriculture and cities as well as forested area. Abandoned land is included in the categories “scrubland” and “other land”. Management intensification is not included per se in HILDA+, but fertiliser and pesticides are able to at least partially capture intensely managed areas. We were unable to represent livestock density and land degradation.

A forest management layer from HILDA+, where all pixels with a fraction of management higher than 50% were classified as managed and all others as unmanaged, was used to determine the level of management in the forest class of the HILDA+ data. All forest pixels of the layer with land use classes were checked against the forest management layer and all unmanaged forest received a score of 0.0, managed forest between 0.1 and 0.5 – depending on the fraction of managed forest in each pixel. Deforestation (as part of resource extraction) is partially represented, because not all forest replacements in the HILDA+ data can be traced back to deforestation and logging (they could be the result of wildfires etc.). We included the forest management layer, because it was a good way to distinguish between “natural” forest and anthropogenically influenced forest and thus minimise erroneous pressure scores for forests.

Land use transitions were taken as a change in land use in a pixel from one year to the next. For the BPI, five years were summarised, i.e. the number of transitions within five years before the respective year; the assumption being that more frequent change leads to more disturbance for biodiversity, since it implies vegetation and habitat change or clearance (Jung et al., 2019). This layer is different from the land use classes layer in that here we are only interested in the frequency of change rather than the land use types involved.

### 4.3 Mining

Availability of spatio-temporal data on resource extraction is limited and we are not aware of a gridded continuous timeseries on mining activities. For our purpose, we chose a gridded dataset that combines data on mining activities from different points in time between 2000 and 2017 into a single dataset for one point in time at a 5 arc-minute resolution (Maus et al., 2020). In our view, these data reflect the direct impacts of mines on biodiversity better than country-level data of mining exports. The data we use includes information on open cuts, tailing dams, waste rock dumps, water ponds and processing infrastructure, and shows the size of the mining area per pixel, i.e., the higher the value, the more area within that pixel is affected by mining. Mining is our only stable variable within the BPI and it represents the non-living materials part of the resource extraction driver defined by IPBES.

### 4.4 Wildlife trade (exports)

To represent the extraction of living materials, we included wildlife trade data, which is based on the CITES (Convention on International Trade in Endangered Species) database and has been published in Jackson et al. (2023). The data contain a list of the number of wild species exported from a country (limited to countries reporting to CITES) within each year from 2000-2020. As with the trade data (see below), we performed back casting in the form of regression to add the years 1990-1999. This back casting was performed on the basis of the entire dataset to avoid undue influence of the rapidly-decreasing trade values in the first few years after 2000. The data were then rasterised using a shapefile. Each pixel within one country received the same value.

### 4.5 Precipitation and temperature

The WMO (World Meteorological Organization) uses seven different climate indicators to quantify climate across time and space, including surface temperature and precipitation (Climate indicators, 2022). The data used were ERA5 land, with a spatial resolution of 10 km and an annual availability from 1950-2021. The data are provided as monthly means and therefore first had to be averaged to annual data. Then the 30-year mean between 1951 and 1980 could be calculated. For the BPI, temperature and precipitation anomalies were calculated by subtracting the 5-year mean of each year and the previous four years (e.g., 1995-2000 for the year 2000) from the 30-year reference mean from 1951-1980. Since we considered all deviations, i.e., hotter or colder and drier or wetter years respectively, from the long-term mean as a negative impact on biodiversity, we took the absolute values of the deviations for further analysis. However, whilst precipitation had positive and negative deviations (i.e. drier or wetter conditions than usual) global temperature anomalies only showed positive deviations from the long-term mean (i.e. all regions showed increases in or stable temperature). With the mentioned variables we are able to include temperature increase and changes in precipitation - two important aspects within the climate change driver – in the BPI. We did not include any data showing sea level rise and ocean acidification, since the marine realm was not part of this analysis.

### 4.6 Pesticides

For the pesticides data we combined yearly national statistics on the agricultural use of pesticides (all pesticides, including insecticides, herbicides, fungicides, plant growth regulators, rodenticides) in tonnes per year from the FAO (Pesticides Use, 2023) with the cropland area derived from HILDA+, resulting in a dataset showing the pesticide use in tonnes per year on each cropland grid cell at 1 km resolution. This meant that for each country, all cropland pixels were assumed to have the same amount of pesticide use. Pesticides and fertiliser (below) are primarily pollutants from agriculture; we do not represent pollution from other sources (e.g. air pollution, microplastics etc.). Pesticides as well as fertilisers are used as application in tonnes/kilograms, meaning that we do not have values for dissolved or emitted contaminants. The BPI also lacks data for solid waste for which no suitable datasets were found.

### 4.7 Fertiliser

The fertiliser use data came from Tian et al. (Tian et al., 2022), which includes spatial information on ammonium, nitrate and manure inputs to pasture and cropland at a 5 arc-minute resolution. The data were constructed, amongst others, from FAO statistics and the HYDE3.2 land use data. We summed all three variables since we were only interested in the total amount of fertiliser applied. The data were only available until 2019, hence the values from 2019 were also used for 2020. We note the potential for inconsistency here with the HILDA+ data used above, but also the benefit of capturing uncertainty in cropland extent between two major land use reconstruction data products.

### 4.8 Nighttime lights

The often-used data set DMSP NTL (Defense Meteorological Satellite Program nighttime light) was harmonized (because data after 2013 were not available) with a new data set from the VIIRS (Visible Infrared Imaging Radiometer Suite) satellite and combined into a new data set showing DN (digital number) values from 0-63. This has a spatial resolution of 1 km and an annual data availability from 1992 to 2018. (Li et al., 2020). To avoid spurious and uncertain lights (such as auroras), we deleted all DN values equal to or below 23. Here again, we extended the dataset to the missing years. Values from 1992 were used for 1990-1991 and values from 2018 for 2019-2020, based on the assumption that maintaining pressures over short periods was more reasonable than creating arbitrary variation

### 4.9 Trade (imports)

Global data on trade in a gridded format does not exist. A dataset from the Chatham House Resource Trade Database (CHRTD) shows the annual imports and exports of all countries worldwide in weight and value of goods from 2000-2020. Because these data are at national scale and do not cover the first decade of our analysis, we extrapolated the dataset back to 1990. We did so by finding the best fitting line to the data from 2000 onwards and then used that linear model to back cast to 1990. This tabular dataset was merged with a shapefile containing country borders and then rasterised, i.e. a gridded time series was calculated. We use trade as a proxy for the potential invasion by non-native species, with the trade data reflecting a value of potential threat from invasive species, not actual invasion, in each cell per country. The potential threat is derived from the quantity of imported goods. Trade has proved to be a good approximation for alien (invasive) species and is widely seen as the main reason for the distribution of alien species (Balvanera et al., 2019; Hulme, 2009; Roy et al., 2024; Seebens et al., 2017). Connectivity was not used as it is so dependent on the species concerned, and so would be unable to provide a general proxy for invasion potential.

### 4.10 Biodiversity Pressure Index maps

We resampled all datasets to a resolution of 0.1° x 0.1°. In order to be able to integrate the datasets into a single index, we classified the input data on a scale from 0 to 1 following a Minimum-Maximum-Normalisation approach. The sum of the resulting standardised pressure layers made up the terrestrial BPI. The BPI itself was normalised again so that the range of the BPI per pixel ranged between minimum 0 and maximum 1, depending on the number and severity of the pressures. We assumed the higher the values of any input variable the higher the stress on biodiversity. We did not, however, give any weighting to the variables as we wanted to represent the actual human pressure within each grid cell independently from the effects on biodiversity. In effect this means there is an equal weighting between the variables, but in the absence of further evidence there is no basis on which to weight the variables unequally. Furthermore, the effect of these pressures on biodiversity is likely to be variable (depending e.g. on local habitats, species compositions and the species itself) and needs targeted analysis. However, the effects of the individual drivers can still be examined independently as shown in Figs. 3 and 4.

All input variables and therefore the resulting BPI are skewed to the right – meaning that there are a large number of small values and few very high numbers. We therefore constructed an alternate quantification of the BPI using percentile ranking of each input dataset to show where pressures varied in relative, rather than absolute, terms. To determine a threshold for ‘no or little pressure’, we followed the approach of Williams et al., who determined a threshold (4 out of 50 pressure points) according to findings regarding species extinctions risk and ecosystem alterations. Areas with a value greater than the threshold are no longer considered natural, but human-dominated (Williams et al., 2020). This led to a threshold of less than 4 in the HF maps (equal to less than 8% of the pressure values). We decided on a threshold of 0.05 in the BPI (corresponding to 5% of the pressure values), because this also matches the point where values become skewed.

### 4.11 Analysis

To calculate the change in area where BPI met a certain condition, or to analyse global and regional means and changes, we applied an area-weighting factor to the BPI.

Difference maps were calculated by subtracting the baseline of the BPI, i.e., 1990, from each year element-wise (per pixel). The result shows areas with increases or decreases in BPI over 30 years and the magnitude of the change.

For the regional analysis, we used data on world regions from the World Bank (World Bank Group, 2017) and data on biome extents from IPBES (Niamir and Obura, 2020) to calculate summaries of BPI for different world regions and biomes, respectively.

The most dominant pressure in each BPI pixel was calculated by ordering the individual pressures by absolute magnitude and then extracting the highest pressure with a difference to the second highest pressure of at least 5%. If no highest pressure was detectable then the pixel was classified as having no dominant pressure.

## Code and data availability

The python and R codes used in this study can be obtained from the authors upon request.

The datasets used to calculate the BPI are openly available (see links in SI). Data of the absolute and relative BPI can be found and freely downloaded at Pangaea repository (https://doi.org/10.1594/PANGAEA.969567).

## Supplement link

The link to the supplement will be included by Copernicus, if applicable.

## Author contributions

KR, CB, AA and MR designed the research project, including the methods used. KR carried out the data search, preprocessing and analysis. KR wrote the manuscript with contributions from all co-authors.

## Disclaimer

Copernicus Publications adds a standard disclaimer: “Copernicus Publications remains neutral with regard to jurisdictional claims made in the text, published maps, institutional affiliations, or any other geographical representation in this paper. While Copernicus Publications makes every effort to include appropriate place names, the final responsibility lies with the authors. Views expressed in the text are those of the authors and do not necessarily reflect the views of the publisher.”

Please feel free to add disclaimer text at your choice, if applicable.

## Acknowledgements

We thank Karina Winkler, Joanna Raymond and Christof Lorenz for their support during the data processing.

## Review statement

The review statement will be added by Copernicus Publications listing the handling editor as well as all contributing referees according to their status anonymous or identified.

