## Supplementary information for "Expansion and increase of human pressures on global land ecosystems between 1990 and 2020" (from another version of this preprint: version 1, 2026-04-20)

### Content

#### Tables

Table S1: Overview of the five IPBES drivers

Table S2: Comparison of indices similar to the BPI

Table S3: List of original input variables

Table S4: Mean absolute and relative BPI in biomes and world regions

#### Figures

Fig. S1: Biome classification from IPBES

Fig. S2: World region classification

Fig. S3: Timeline global mean BPI

Fig. S4: Proportionately highest pressures behind the BPI 1990, 2000, 2010 and 2020

Fig. S5: Percentage change, absolute change and standard deviation of the input variables

Fig. S6: Timelines of individual absolute input variables

Fig. S7: Timelines of absolute individual pressures aggregated to world regions

Fig. S8: Timelines of absolute individual pressures aggregated to biomes

Fig. S9: Changes in pressure (2020 compared to 1990) of individual absolute input variables

Fig. S10: Changes in pressure (2020 compared to 1990) of individual relative input variables

Table S1: Overview of the five IPBES drivers. Big boxes are the overarching topics, left columns reflect the content of the IPBES Global Report, 2019 and right columns show the variables included in the BPI

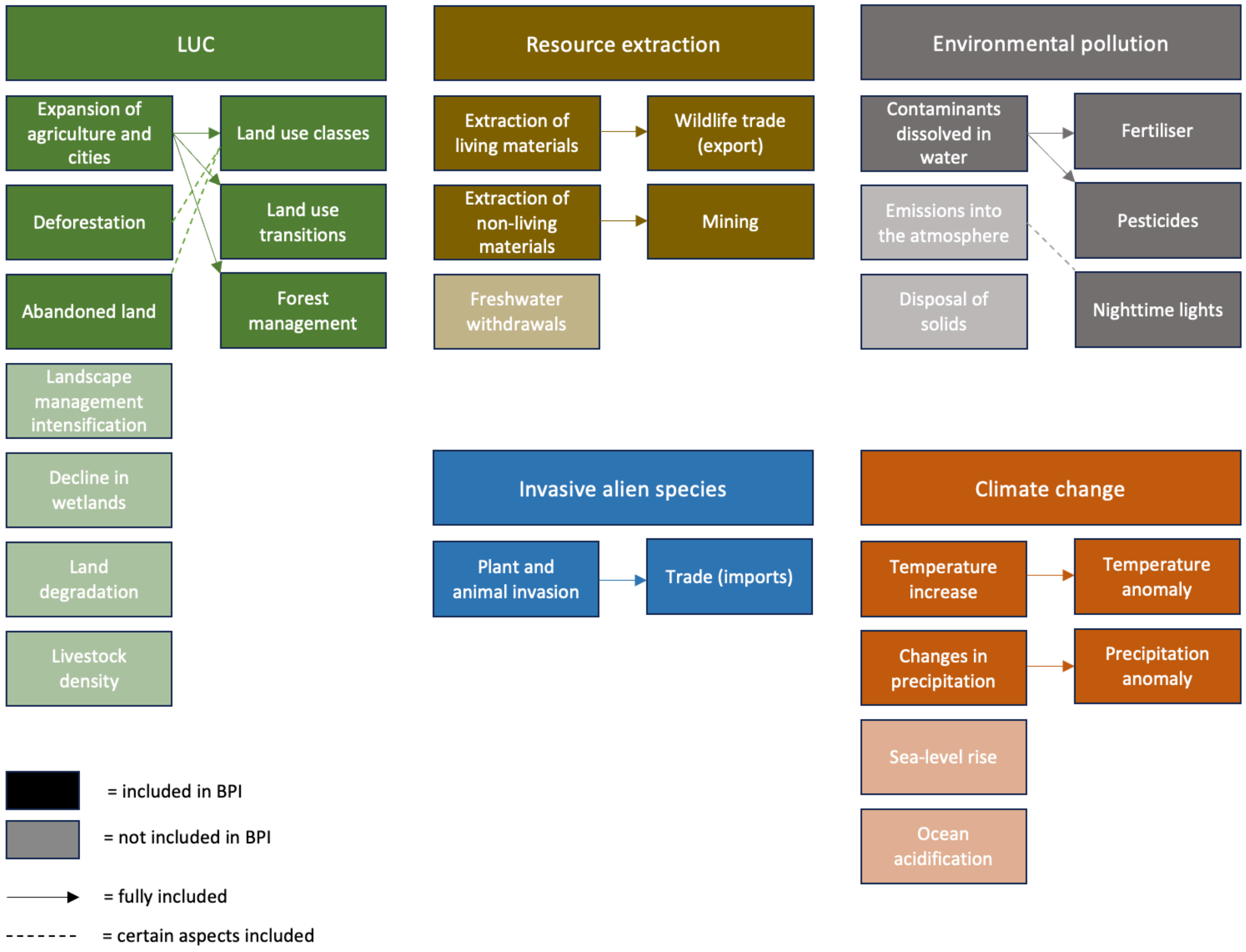

Table S2: Comparison of indices similar to the BPI

| **Index/dataset** | **Variables used** | **Temporal resolution** | **Spatial resolution** | **Shortcomings** | **Link to paper** |
|---|---|---|---|---|---|
| Human Footprint, Last of the wild project | • Population density<br>• Land transformation<br>• Access (roads, railways, waterways)<br>• Electrical power infrastructure (nighttime lights) | One point in time (about 1992) | 1 km | Climate data, pollution data, resource extraction data not included, as well as fewer information on land use. Furthermore, it is only one point in time, so no temporal changes can be analysed. | Sanderson et al., 2002 |
| Temporal Human Pressure Index | • Human population density<br>• Land transformation (HYDE 3.1)<br>• Electrical power infrastructure (nighttime lights) | 1990 and 2010 | 10 km | Climate data, pollution data, resource extraction data not included, as well as fewer information on land use. Furthermore, it contains only 2 years and no complete timeseries. | Geldmann et al., 2014 |
| Human Footprint Maps | • Built environments<br>• Population density<br>• Electric infrastructure (night time lights)<br>• Crop lands (UMD Land Cover and GlobCover)<br>• Pasture lands (UMD Land Cover and GlobCover)<br>• Roads<br>• Railways<br>• Navigable waterways | 1993 and 2009 | 1 km | Climate data, pollution data not included, as well as fewer information on land use. It also only contains 2 years and 3 pressures are static, i.e. the same data is used for both years. | Venter et al., 2016 |
| Human Modification Index | • Urban and built-up areas<br>• Agriculture (ESA CCI, GLS)<br>• Energy production & mining<br>• Transportation corridors | 1990, 2000, 2010, 2015 and 2017 (with additional stressors) | 0.09 $km^2$ (300 x 300 m) | Climate data not included, pollution from agriculture not included. No annual timeseries. And 2017 not comparable to the other years. | Theobald et al., 2020 |

| | | | | | |
|---|---|---|---|---|---|
| | • Biological harvesting<br>• Human intrusions<br>• Natural system modification<br>• Air pollution (NOx) | | | | |
| Anthropogenic threat complexes | • Climate change<br>• Human use (mainly land use)<br>• Human population density<br>• Pollution<br>• Transport infrastructure | One point in time | 100 $km^2$ | Similar data used compared to the BPI, however, much coarser grid and also only representing one point in time. | Bowler et al., 2020 |
| Human Footprint | • Built environments<br>• Population density<br>• Electric infrastructure (nighttime lights)<br>• Crop lands (ESA CCI)<br>• Pasture lands (ESA CCI)<br>• Roads<br>• Railways<br>• Navigable waterways | 2000-2018, annual | 1 km | Climate data and pollution data is not included. It covers only 18 years. | Mu et al., 2022 |
| Biodiversity Pressure Index | • Land use (and land use transitions) (HILDA+)<br>• Mining<br>• Wildlife trade (exports)<br>• Pesticides use<br>• Fertiliser use<br>• Night time lights<br>• Temperature and precipitation anomalies<br>• Trade (imports) | 1990-2020, annual | 0.1° (~10 km) | Mining used as a static variable and (wildlife) trade data only available on a national level. | |

Table S3: List of original input variables

| IPBES category | Pressure in BPI | Content used in BPI | Dataset | Spatial resolution | Temporal availability | Source/author and reference | Overview & notes |
|---|---|---|---|---|---|---|---|
| Land use | Land use | Classes: Urban areas, crops, pasture/rangeland, forest, sparse/no vegetation, water, no data | HILDA+ 2.0 | 1 km | 1960-2020 | Winkler et al. (2021) https://doi.pangaea.de/10.1594/PANGAEA.921846 and Winkler et al. unpublished | Land use data chosen with implications for other data which are based on land use data in some way |
| | Forest management | Binary: managed or unmanaged forest | HILDA+ 2.0 | 1 km | 1960-2020 | Winkler et al. unpublished | Only differentiates between managed and unmanaged forest, so no intensity of management |
| | Land use change | Number of land use transitions within 5 years | HILDA+ 2.0 | 1 km | 1960-2020 | Winkler et al. (2021) https://doi.pangaea.de/10.1594/PANGAEA.921846 and Winkler et al. unpublished | Data shows transitions from one year to next |
| Resource extraction | Mining | Fractions of mining within each pixel | Global-scale mining polygons (Version 2) | 5 arc-minute | Combination of data with mining reports between 2000 and 2017 | Maus et al. (2020) https://doi.pangaea.de/10.1594/PANGAEA.942325 | No time series → data was only used as a stable variable for BPI maps |
| | Wildlife trade | Traded species per country per year | Global data based on CITES | National level | 2000-2020 | https://www.sciencedirect.com/science/article/pii/S2351989423003773 | Includes endangered species only and is at national level only |
| Environmental pollution | Fertiliser | Fertiliser application in tonnes/ha/yr | HaNi | 5 arc-minute | 1860-2019 | Tian et al. (2022) https://doi.pangaea.de/10.1594/PANGAEA.942069 | Different nitrogen inputs (no phosphorous): ammonium, nitrate, manure |

| | | | | | | | |
|---|---|---|---|---|---|---|---|
| | Pesticides | Pesticides application on HILDA+ cropland area in tonnes | Pesticides use (total) | Country level | 1990-present | FAOSTAT https://www.fao.org/faostat/en/#data/RP | National statistics of fertiliser input in tonnes. We used a crop mask from HILDA+ to spread fertiliser values equally on all crop pixels within each country → results in potential over- or underestimation of fertiliser per crop pixel |
| | Night time lights | Digital numbers (DN): the higher the value the more light | Harmonisation of DMSP and VIIRS nighttime light data | 30 arc-seconds | 1992-2018 | Li et al. (2020) Link to dataset | Spurious lights within the lower DN values and potentially auroras captured in the northern latitudes → in order to minimize errors, we deleted all values equal or smaller than 23, which means we lost information on rural lights and settlements |
| Invasive species | Trade (imports) | Imports of xxx per country | CHRTD | Country level | 2000-present | Chatham House https://resourcetrade.earth | Trade can only reflect the potential for invasive species and is not an actual count or measurement of invasives |
| Climate change | Precipitation anomaly | Anomaly: positive and negative deviation of precipitation from long-term mean | ERA5-Land monthly averaged data | ~ 9 km | 1950-present | ECMWF https://cds.climate.copernicus.eu/cdsapp#!/dataset/reanalysis-era5-land-monthly-means?tab=overview | |
| | Temperature anomaly | Anomaly: positive deviation of temperature from long-term mean | ERA5-Land monthly averaged data | ~ 9 km | 1950-present | ECMWF https://cds.climate.copernicus.eu/cdsapp#!/dataset/reanalysis-era5-land-monthly-means?tab=overview | |

Table S4: Mean absolute and relative BPI in biomes and world regions (1990 and 2020)

| **Biome** | **Absolute BPI** | | **Relative BPI** | |
|---|---|---|---|---|
| | **1990** | **2020** | **1990** | **2020** |
| Tropical and subtropical dry and humid forests | 0.11 | 0.18 | 0. 36 | 0.40 |
| Temperate and boreal forests and woodlands | 0.09 | 0.12 | 0.27 | 0.27 |
| Mediterranean forest, woodlands and shrubs | 0.13 | 0.17 | 0.39 | 0.39 |
| Tundra and high mountain habitats | 0.04 | 0.09 | 0.16 | 0.18 |
| Tropical and subtropical savannas and grasslands | 0.11 | 0.14 | 0.32 | 0.34 |
| Temperate grasslands | 0.14 | 0.17 | 0.35 | 0.34 |
| Deserts and xeric shrublands | 0.06 | 0.11 | 0.22 | 0.22 |
| Wetlands | 0.16 | 0.22 | 0.36 | 0.41 |
| Cryosphere | 0.01 | 0.01 | 0.03 | 0.02 |
| Global | 0.06 | 0.09 | 0.20 | 0.20 |

| **World region** | **Absolute BPI** | | **Relative BPI** | |
|---|---|---|---|---|
| | **1990** | **2020** | **1990** | **2020** |
| East Asia & Pacific | 0.11 | 0.24 | 0.33 | 0.34 |
| Europe & Central Asia | 0.06 | 0.08 | 0.21 | 0.21 |
| Latin America & Caribbean | 0.10 | 0.12 | 0.32 | 0.36 |
| Middle East & North Africa | 0.05 | 0.09 | 0.20 | 0.21 |
| North America | 0.09 | 0.10 | 0.25 | 0.24 |
| South Asia | 0.15 | 0.22 | 0.40 | 0.42 |
| Sub-Saharan Africa | 0.09 | 0.13 | 0.29 | 0.31 |
| Global | 0.09 | 0.14 | 0.28 | 0.30 |

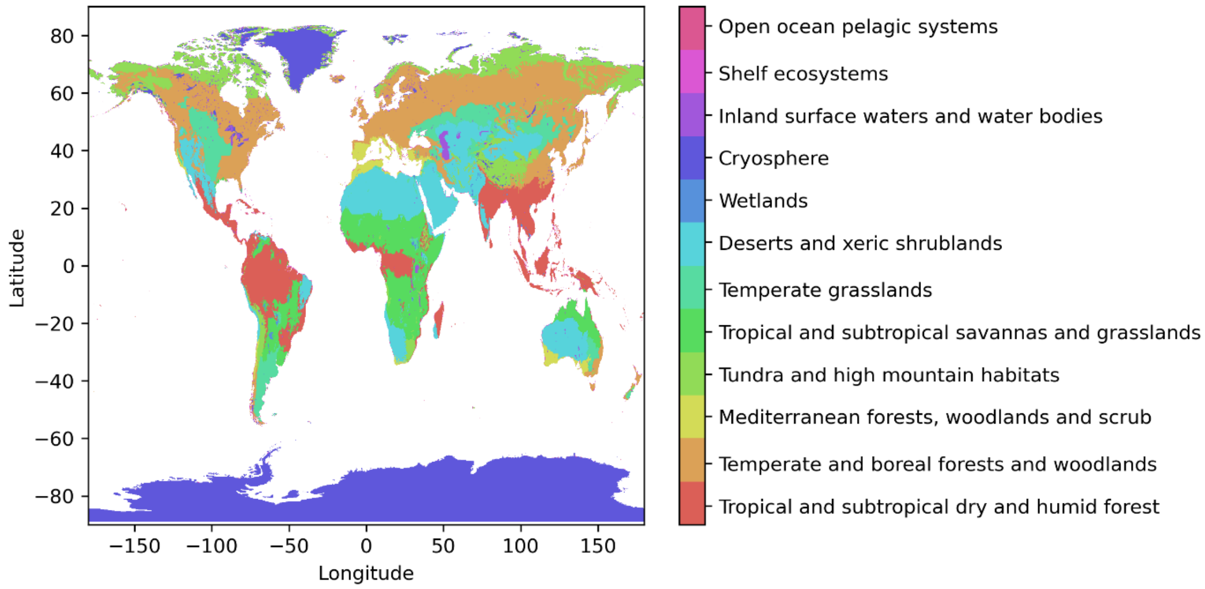


Fig. S1: Biome classification from IPBES following (Niamir & Obura, 2020) https://zenodo.org/records/3975694

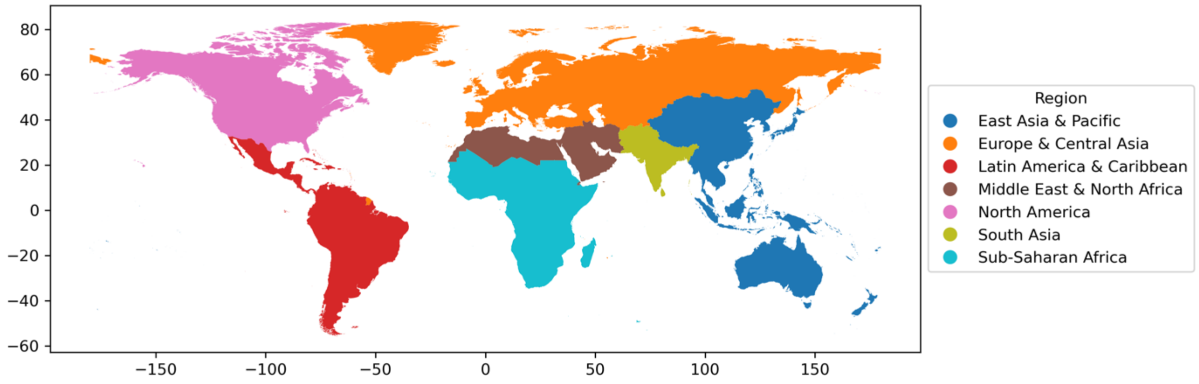


Fig. S2: World region classification following (World Bank Group, 2017) https://ourworldindata.org/grapher/world-regions-according-to-the-world-bank

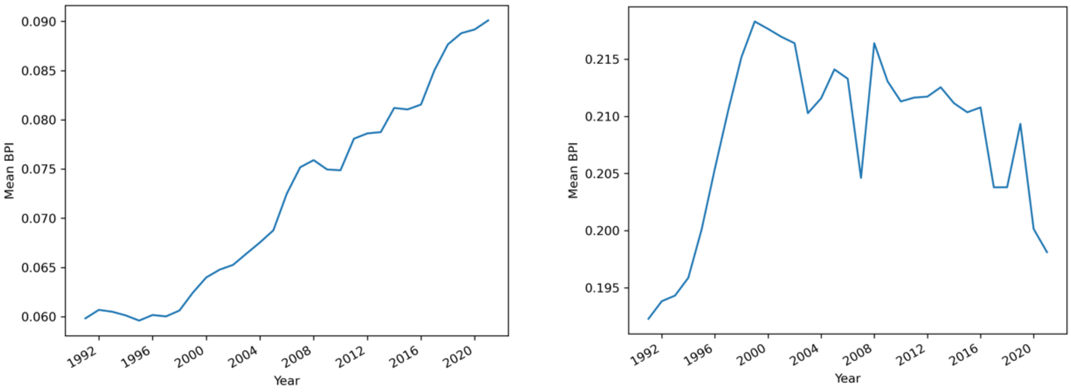


Fig. S3: Timeline global mean BPI (absolute values left, relative values right) from equal area projection

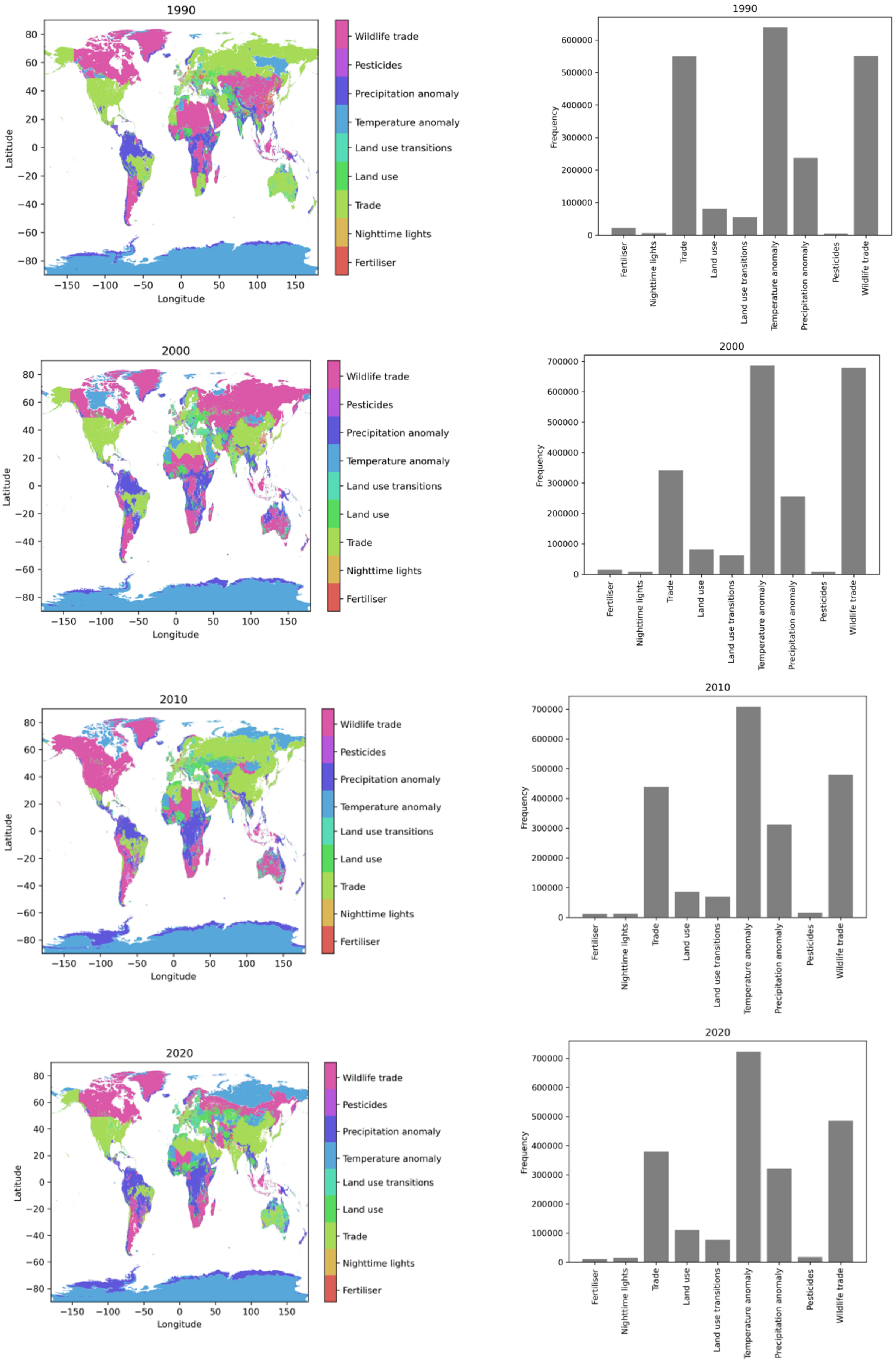


Fig. S4: Proportionately highest pressures behind the BPI 1990, 2000, 2010 and 2020

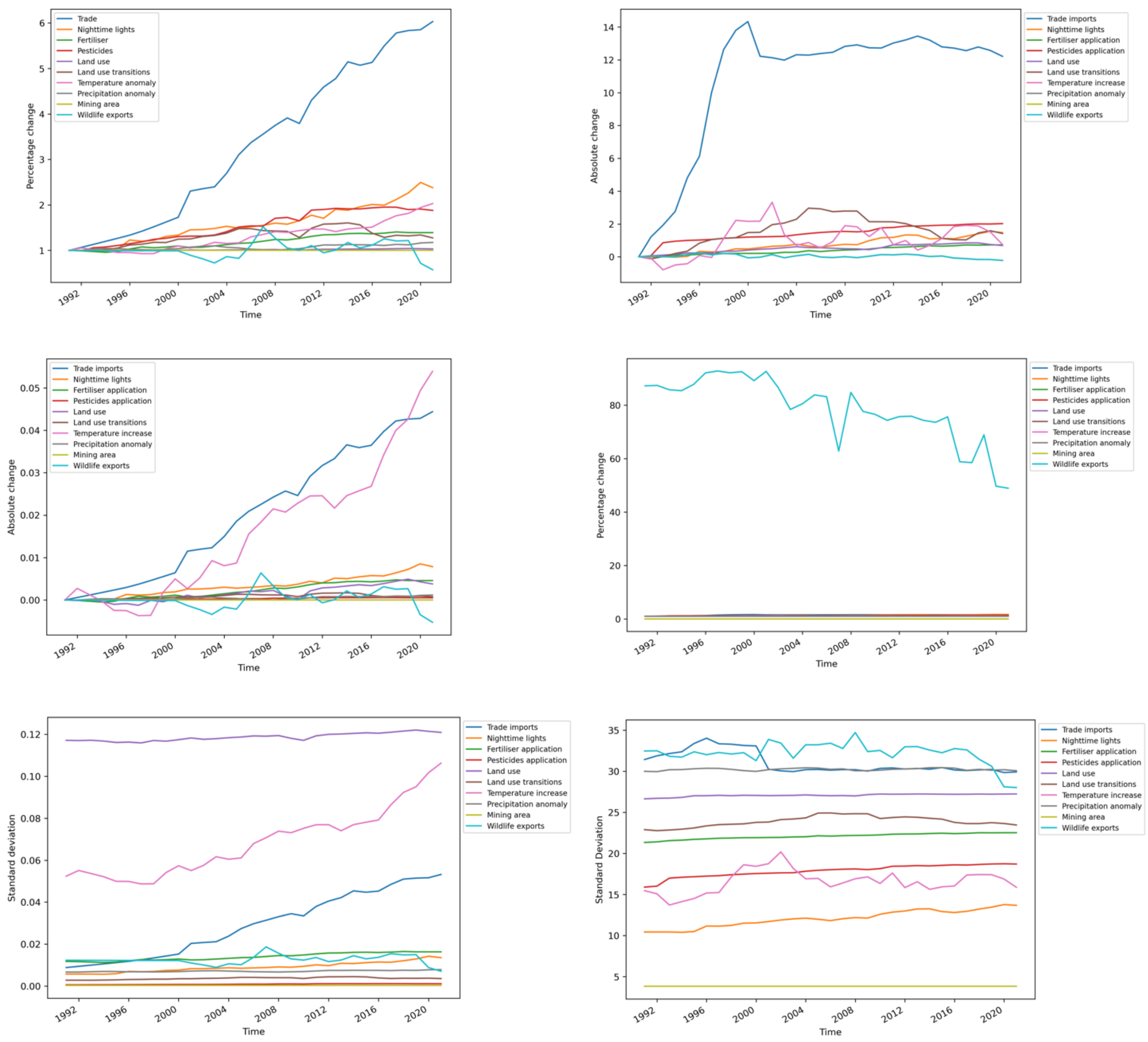


Fig. S5: Percentage change (a), absolute change (b) and standard deviation (c) of the input variables to the BPI of the absolute (left column) and relative (right column) inputs (legend valid for all plots)

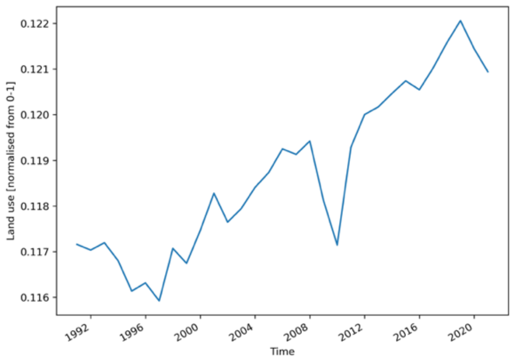
Land use [normalised from 0-1]
Time

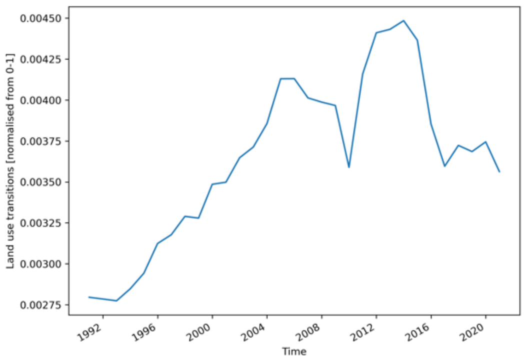
Land use transitions [normalised from 0-1]
Time

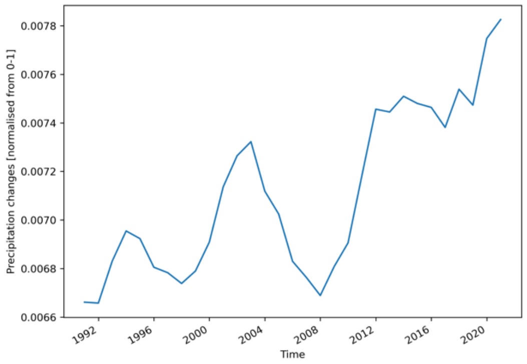
Precipitation changes [normalised from 0-1]
Time

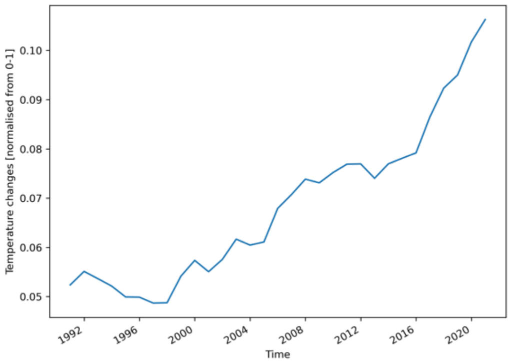
Temperature changes [normalised from 0-1]
Time

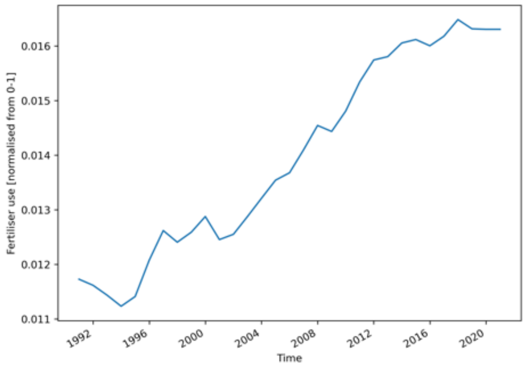
Fertiliser use [normalised from 0-1]
Time

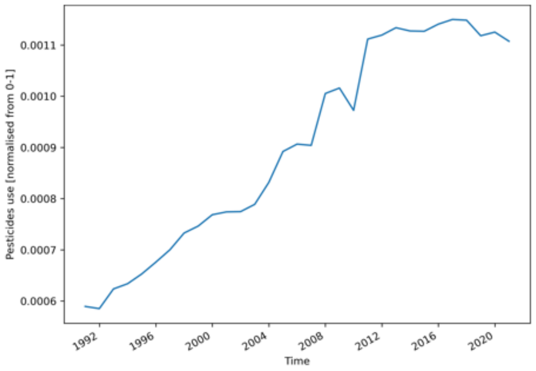
Pesticides use [normalised from 0-1]
Time

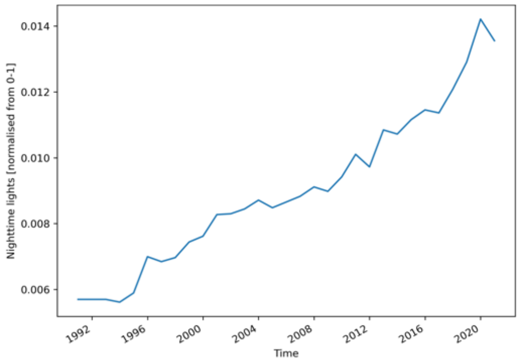
Nighttime lights [normalised from 0-1]
Time

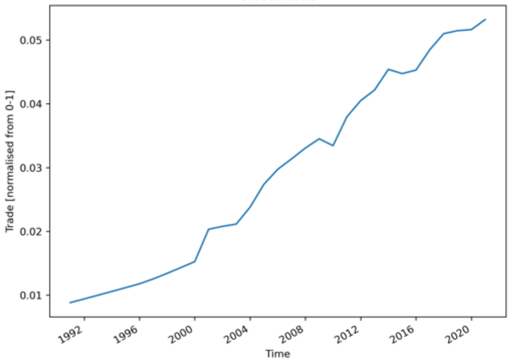
Trade [normalised from 0-1]
Time

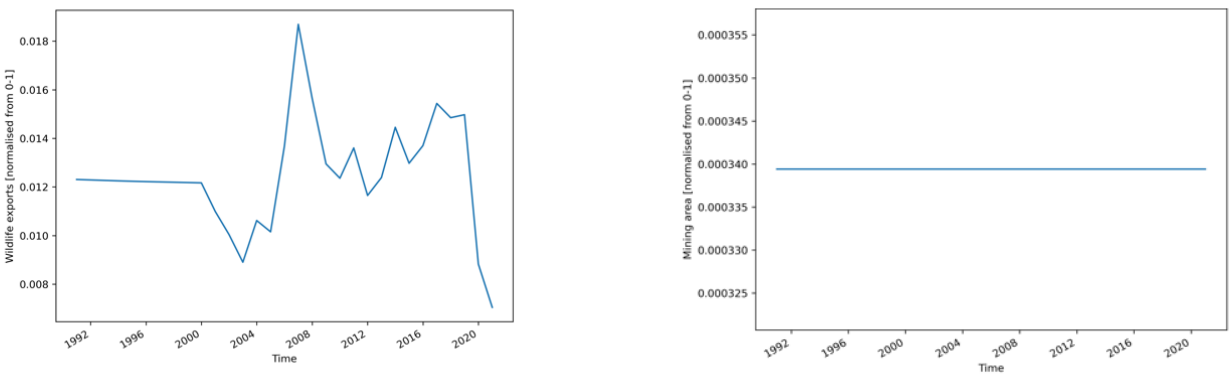


Fig. S6: Timelines of individual absolute input variables to the BPI 1990-2020

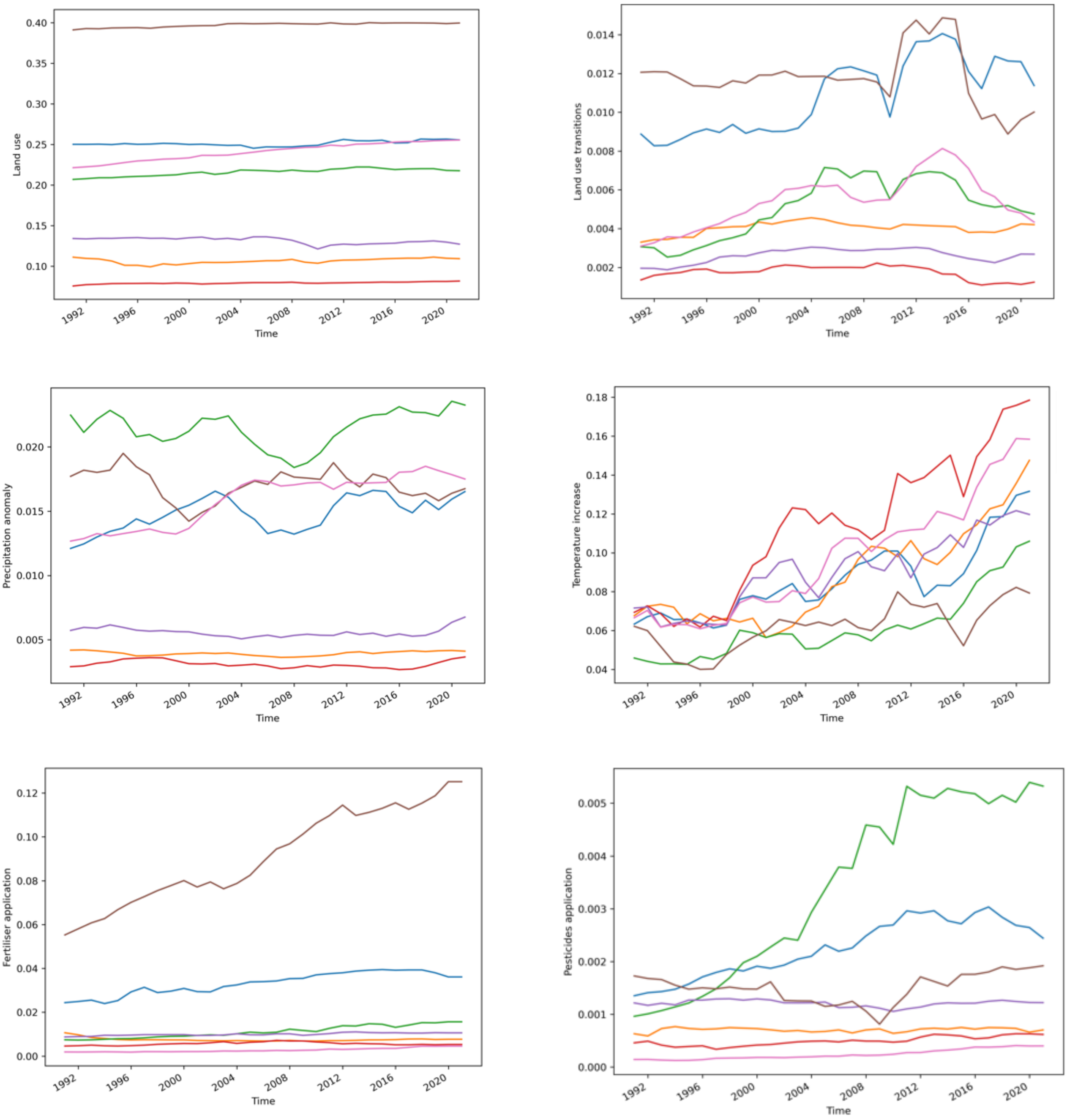

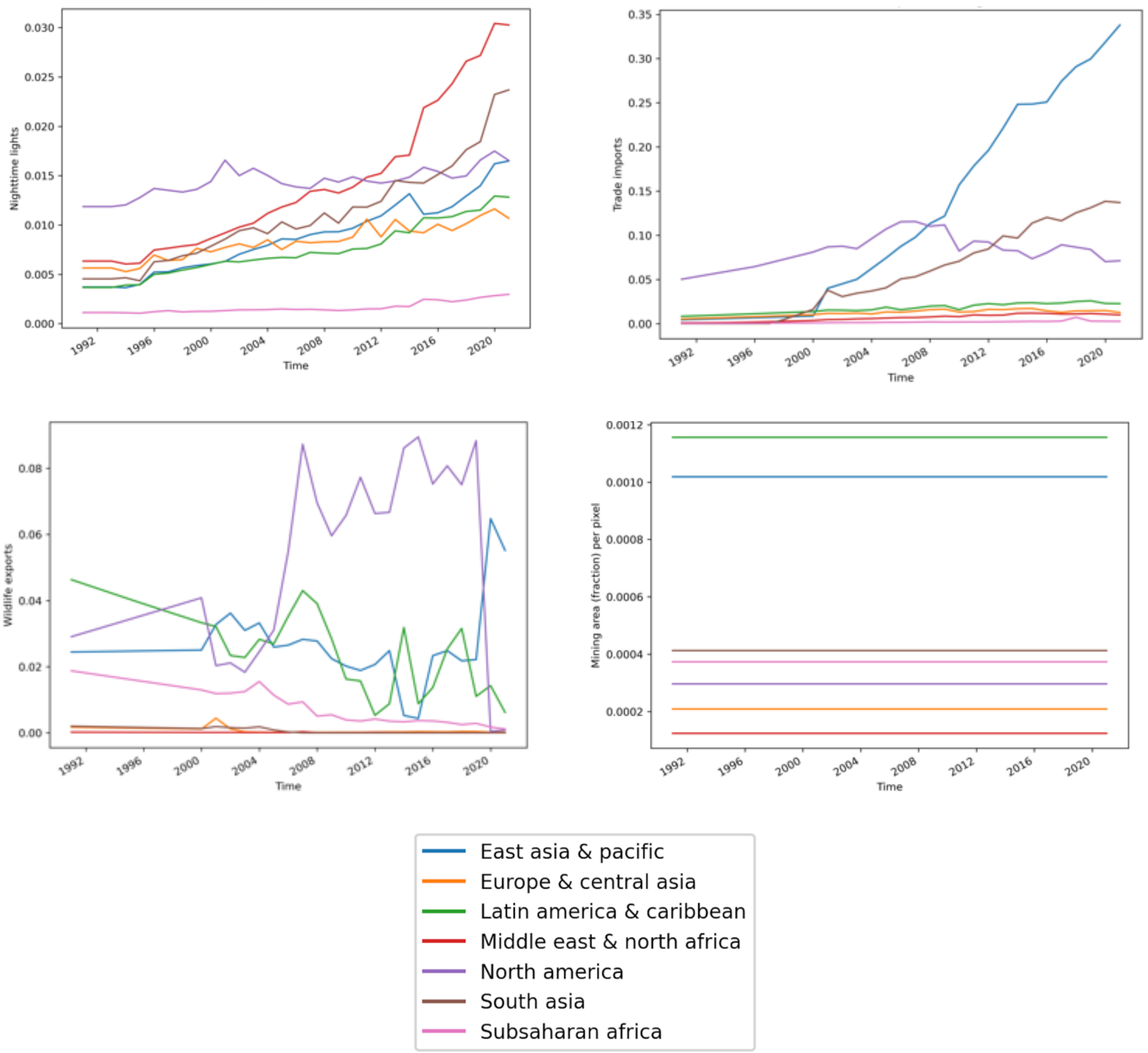


Fig. S7: Timelines of absolute individual pressures aggregated to world regions (legend valid for all plots)

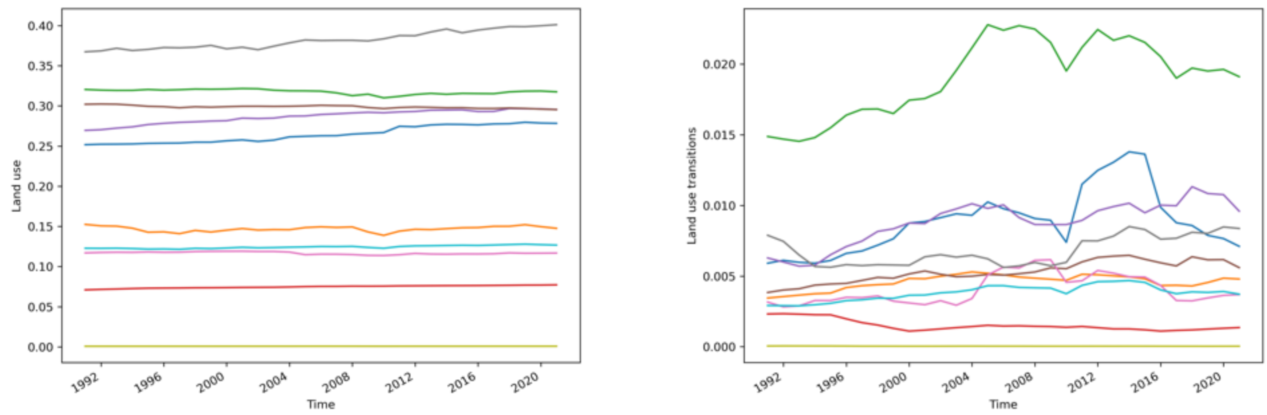

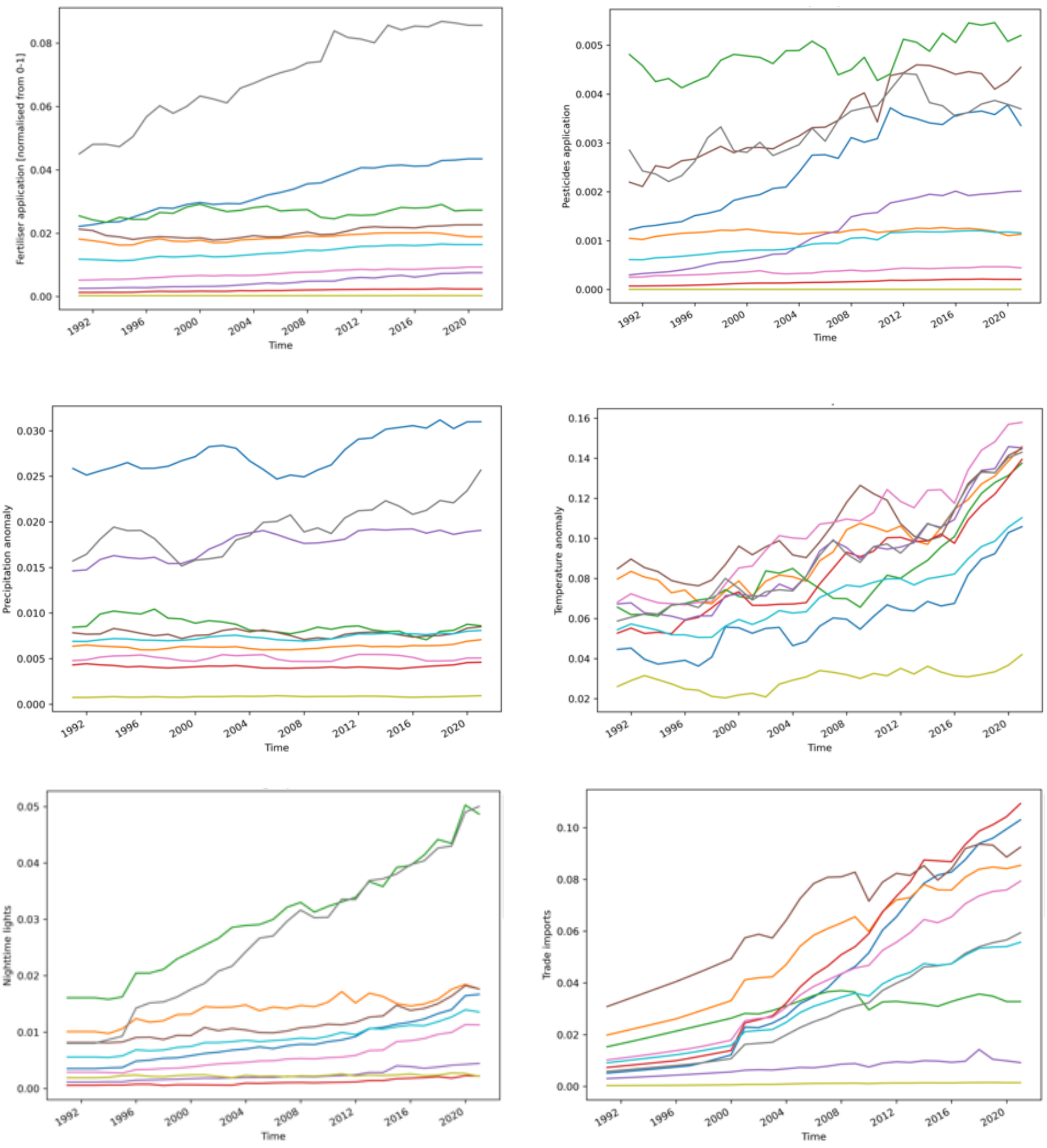

Fertiliser application [normalised from 0-1]
Time
Pesticides application
Time
Precipitation anomaly
Time
Temperature anomaly
Time
Nighttime lights
Time
Trade imports
Time

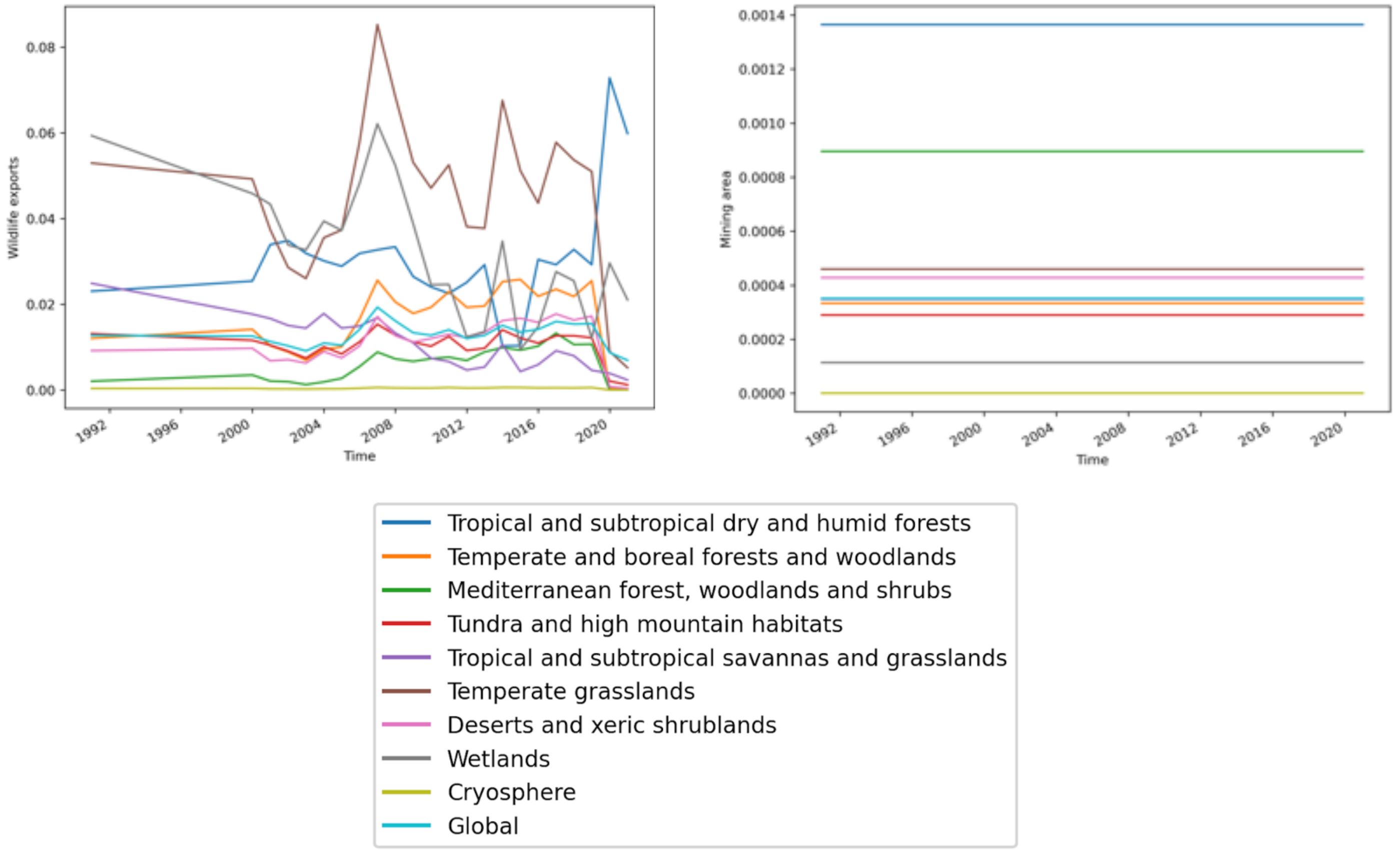


Fig. S8: Timelines of absolute individual pressures aggregated to biomes (legend valid for all plots)

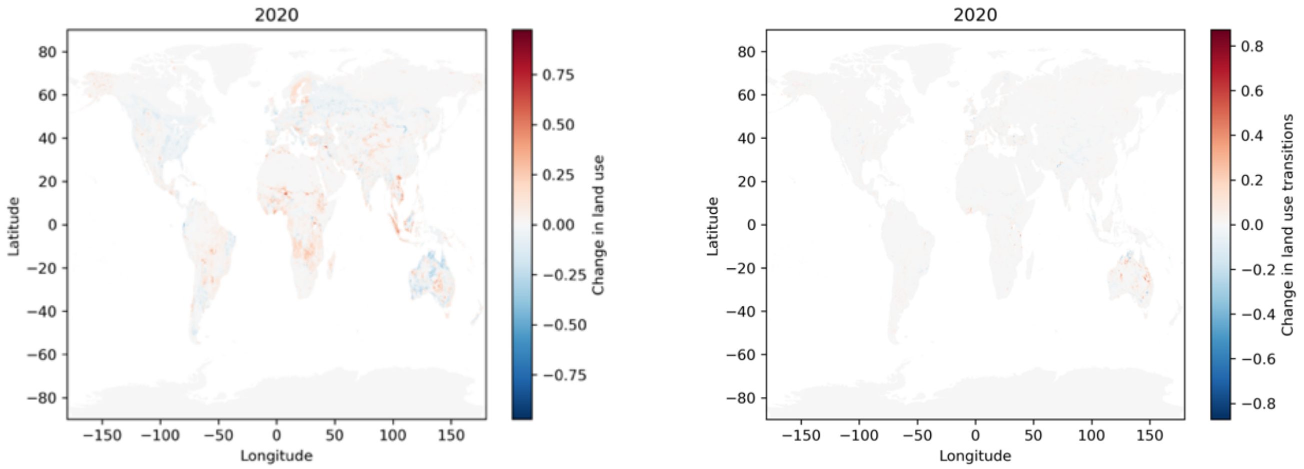

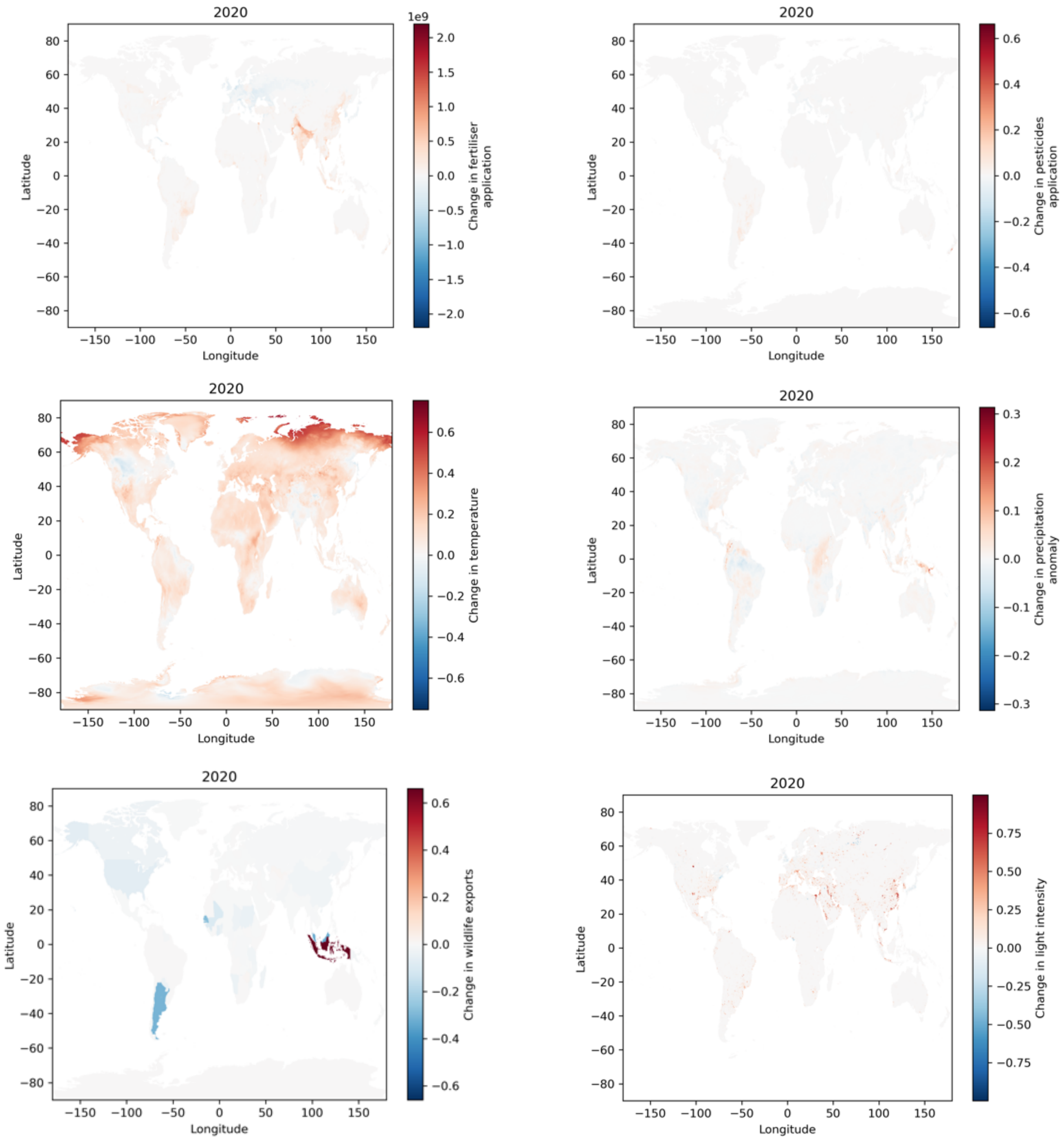


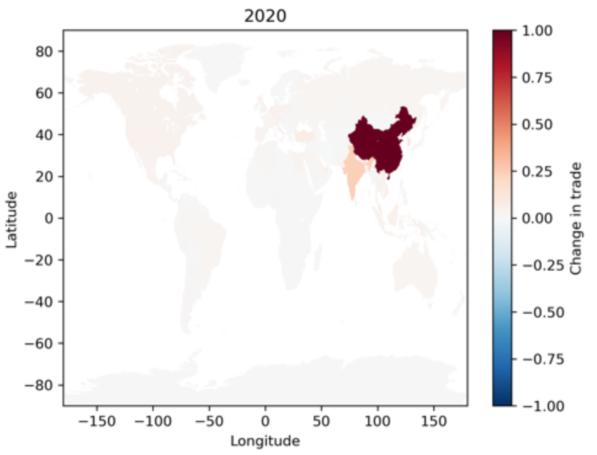


Fig. S9: Changes in pressure (2020 compared to 1990) of individual absolute input variables

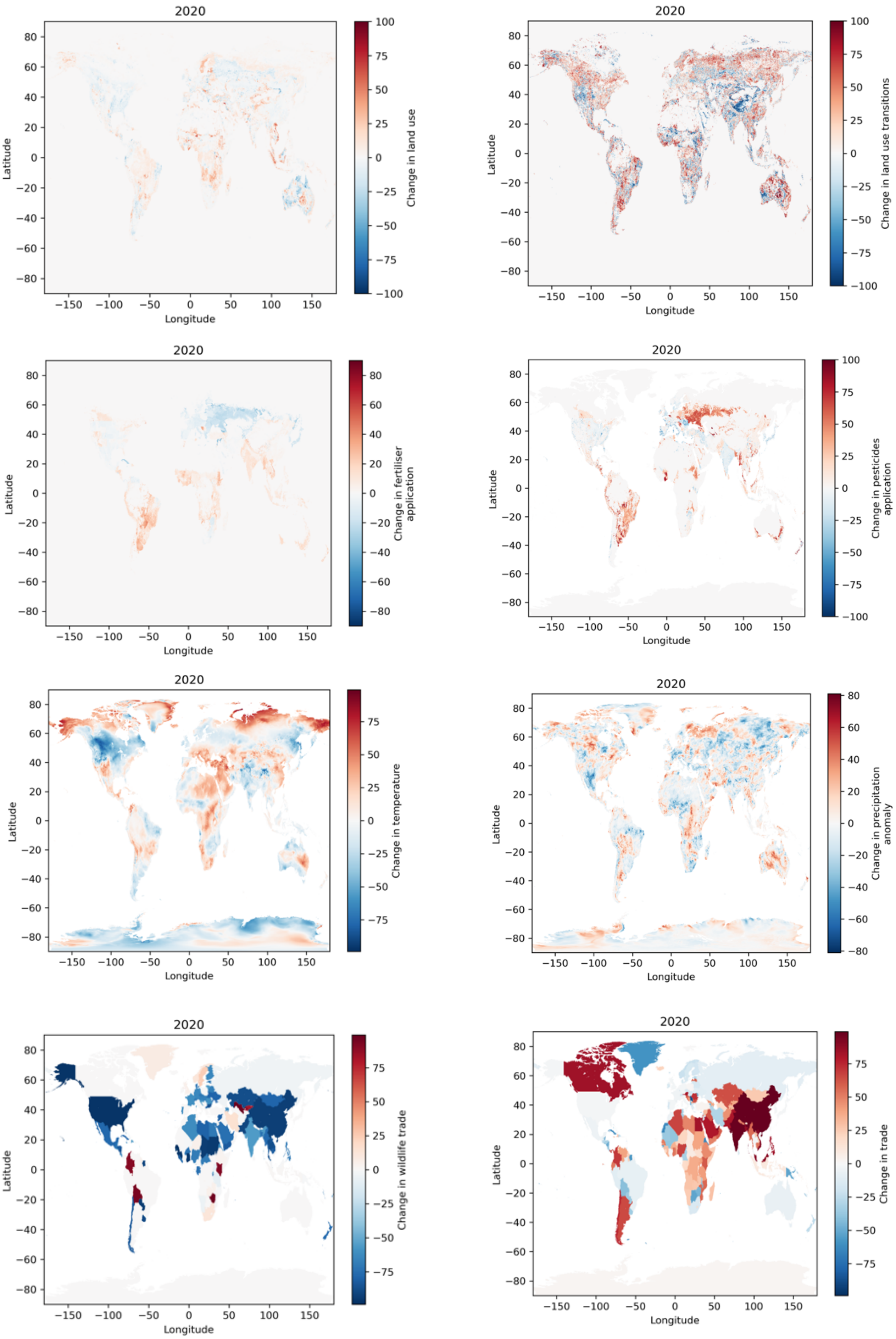

2020
Latitude
Longitude
Change in land use
2020
Latitude
Longitude
Change in land use transitions
2020
Latitude
Longitude
Change in fertiliser application
2020
Latitude
Longitude
Change in pesticides application
2020
Latitude
Longitude
Change in temperature
2020
Latitude
Longitude
Change in precipitation anomaly
2020
Latitude
Longitude
Change in wildlife trade
2020
Latitude
Longitude
Change in trade

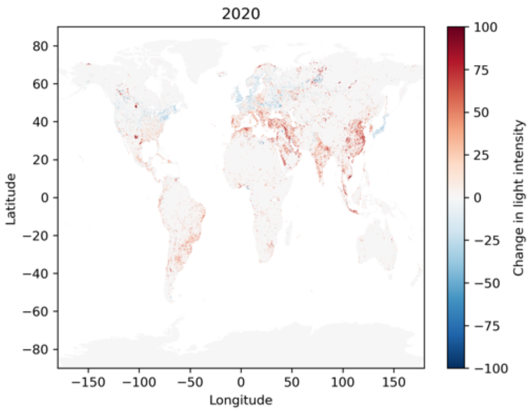


Fig. S10: Changes in pressure (2020 compared to 1990) of individual relative input variables
